# Experimental procedures for flow cytometry of wild type mouse brain: A systematic review

**DOI:** 10.1101/2023.07.06.547972

**Authors:** Robert C. Sharp, Dylan T. Guenther, Matthew J. Farrer

## Abstract

**Objective:** To systematically review the neuroimmunology literature to determine the average immune cell counts reported by flow cytometry in wild-type (WT) homogenized mouse brain.

**Background:** Mouse models of gene dysfunction are widely used to study age-associated neurodegenerative disorders, including Alzheimer’s disease and Parkinson’s disease. The importance of the neuroimmune system in these multifactorial disorders has become increasingly evident, and methods to quantify resident and infiltrating immune cells in brain, including flow cytometry, are necessary. However, there appears to be no consensus on the best approach to perform flow cytometry or quantify/report immune cell counts. The development of more standardized methods would accelerate neuroimmune discovery/validation by meta-analysis.

**Methods:** Examination of the PROSPERO registry confirmed a systematic review of ‘neuroimmunology’ by ‘flow cytometry’ has yet to be reported. A protocol for a systematic review was subsequently based on the Preferred Reporting Items for Systematic Reviews and Meta-Analyses (PRISMA) using Studies, Data, Methods and Outcomes (SDMO) criteria. Literature searches were conducted in Google Scholar and PubMed databases. From that search, 900 candidate studies were identified and 437 studies were assessed for eligibility based on formal exclusion criteria.

**Results:** Out of the 437 studies reviewed, 58 were eligible for inclusion and comparative analysis. Each study assessed immune cell subsets within homogenized mouse brain and used flow cytometry. Nonetheless, there was considerable variability in methods, data analysis, reporting and results. Descriptive statistics are presented on study designs and results, including medians with interquartile ranges (IQRs) and overall means with standard deviations (SD) for specific immune cell counts and their relative proportions, within and between studies. Results derived from 58 studies reported the most abundant immune cells within the brains were TMEM119^+^ microglia, bulk CD4^+^ T cells, and bulk CD8^+^ T cells.

**Conclusion:** Experiments to conduct and report flow cytometry data, derived from WT homogenized mouse brains, would benefit from a more standardized approach. While within study comparisons are valid, variability in methods counts of immune cell populations are too broad for meta-analysis. Inclusion of a minimal protocol with more detailed methods, controls and standards could enable this nascent field to compare results across studies.

## 1 Introduction

The mouse has been used to model neurological disorders for many decades, whether lesion models (i.e. inducing a stroke in a mouse specimen), using transgenic or more physiologic gene knock-out or mutant knock-in approaches (1–3). Some illustrations of mouse modeling for neurologic and neurodegenerative disorders include work in Alzheimer’s disease (AD), Parkinson’s disease (PD), Huntington’s disease (HD), multiple sclerosis (MS), psychological/intellectual disabilities (i.e. depression or Down’s syndrome), traumatic brain injuries (TBI) and prion diseases (1–3). Most of these disorders are multifactorial, with genetic and environmental components, for which the immune system may provide some integration. Consequently, there has been growing interest in neuroimmunology, and not just focused on resident immune cells within the brain, but also on infiltrating peripheral immune cells (4–7). Recent studies of gut microbiota have highlighted nervous and immune communication between the gut and central nervous system (CNS) (8–10). With this resurgence, researchers have employed traditional (i.e. immunofluorescence/histology slide staining and Western blotting) and contemporary methods (i.e. single-cell genomics and single-cell sorting/staining via flow cytometry) to better characterize specific immune cell subsets within the body and brain.

Specifically, brain resident immune cells, including microglia and astrocytes, have been comprehensively examined in many mouse models of neurologic disorders (4–10). However, the characterization of other immune cell subsets in peripheral blood, the CNS, and within the brain (both residential and infiltrated) has yet to be fully described (4–10). These immune cell subsets include, but are not limited to, natural killer cells (NK cells), T cells, and B cells (4–10). However, validation and comparison through meta-analysis of immunophenotypes of these mouse models might be enabled if researchers utilize more standardized methods and reporting.

Technologic developments for single-cell isolation and analysis, including single-cell RNAseq, mass cytometry (CYTOF), fluorescence-activated cell sorting (FACS) and multi-color flow cytometry derived immunophenotyping, have illuminated a wide variety of scientific fields (11,12). Of these techniques, FACS and flow cytometry immunophenotyping are most frequently used. Reasons include the ease of use in setting up a flow cytometer or sorter for a variety of applications, the sensitivity of the assay, the specificity of the antibodies used, the potential of those antibodies to be used for both flow cytometry and Western blotting for the same target, and the ability to produce qualitative and quantitative data (12,13).

Nevertheless, flow cytometry has its pitfalls as analytic interpretation of the data depends on the user’s preference for gating and target choices (13). Experimentally, flow cytometry is also dependent on the fluorophores and cytometers used, and variation between these instruments may result in false positives and negatives. Nevertheless, such issues can be circumvented by providing multiple controls, such as unstained, isotype antibody-stained and fluorescence minus one (FMO) controls, more rigorously described methods and standardization of flow cytometry protocols.

FACs and flow cytometry immunophenotyping have been insightful and widely used in basic immunology that other scientific fields, including neurology, have begun to adopt these methods (14–16). However, the use of flow cytometry for detecting cell types within the brain and CNS warrants more standardized protocols and reporting, now that their utility has been demonstrated. Although mouse immune profiles within the brain have been identified by high-dimensional single-cell mapping using techniques like mass cytometry (CYTOF)(17), at the time of writing and despite numerous publications, the expected cell counts and proportions of each immune subset within the brain of a wild type (WT) mouse have not yet been clearly defined by standard FACS and flow cytometry. Consequently, we have performed a systematic review focused on neuroimmunology and the use of flow cytometry to detect immune cells derived from WT homogenized mouse brain. In our results, we summarize the number of immune cells overall and estimate the immune subsets that can be detected via flow cytometry immunophenotyping. Our findings demonstrate a critical need for more standardized methods and reporting, and led to best practice recommendations for future publications.

## 2 Methods

### 2.1 Study design

Study design was informed by prior literature (18–20) and based upon the Preferred Reporting Items for Systematic Reviews and Meta-Analyses (PRISMA) criteria (21) and the Cochrane Handbook of reporting methodology reviews, employing SDMO (Types of Studies, Types of Data, Types of Methods, and Types of Outcome Measures) criteria (22,23). Bias assessment for each individual study selected for systematic review inclusion was also conducted using Systematic Review Centre for Laboratory animal Experimentation (SYRCLE) (24) methodology, and subsequent reporting uses the *robvis* R package and Shiny web app (25).

### 2.2 Search strategy

Google Scholar and PubMed databases were used to identify all studies for this systematic review, published between Jan 2013 and July 2023. The search protocol and study design were also assessed within the National Institute of Health (NIH) PROSPERO registry database, which confirmed a review of this topic has not been previously conducted. For database searches, the following combination of keywords were used to identify eligible studies: “flow cytometry” AND “immune subset name examined in study” AND “mouse brain.” For example, a keyword search containing “flow cytometry CD4 T cells mouse brain” was used. “Immune subset name examined in study” is defined as one of the following immune cell subsets:,” “CD4 T cells,” “CD8 T cells,” “double negative DN T cells,” “regulatory T cells TREG,” “follicular helper T cells TFH,” “T helper 1 T cells TH1,” “T helper 2 T cells TH2,” “T helper 17 T cells TH17,” “naïve T cells,” “naïve CD4 T cells,” “naïve CD8 T cells,” “naïve-like T cells,” “central memory T cells TCM,” “central memory CD4 T cells TCM,” “central memory CD8 T cells TCM,” “effector memory T cells TEM,” “effector memory CD4 T cells TEM,” “ effector memory CD8 T cells TEM,” “effector memory T cells re-expressing CD45RA TEMRA,” “effector memory CD4 T cells re-expressing CD45RA TEMRA,” “effector memory CD8 T cells re-expressing CD45RA TEMRA,” “TEMRA-like T cells,” “natural killer cells NK cells,” “dendritic cells DC,” “B cells,” “monocytes,” “macrophages,” “M1 macrophages,” “M2 macrophages,” “TMEM119 microglia,” or “neutrophils.”

### 2.3 Selection and exclusion criteria

Studies were selected from search results employing the following inclusion criteria: 1) any study performed between 2013-2023; 2) any study that contained flow cytometry data identifying immune cell subsets and counts in mouse brain; 3) any study that reported total cell numbers or total live cell percentages for one or more immune cell subsets, and; 4) any study that had WT mice or a non-treated control (when reporting on transgenic mouse models). Studies were excluded from search results based upon the following criteria: 1) studies performed in 2012 and prior; 2) any study not focused on flow cytometry of homogenized mouse brains; 3) rat studies; 4) human studies, or; 5) studies not reporting controls or the background mouse strain.

### 2.4 Data extraction

For each study, information on the mouse strain used, age, and sex were recorded. In addition, methodological information on mouse perfusion, brain extraction and homogenization were recorded. Flow cytometry methods were recorded when the following was reported: 1) cytometer make and model; 2) software for data collection/analysis; 3) full gating strategy; 4) total cells collected per sample; 5) total live cell counts; 6) total immune cell subset percentages calculated directly from live cell counts, and; 7) methods for determining cell counts and/or mean fluorescence intensity (MFI) readings.

Studies were subsequently examined for immune cell subset counts from WT/control mouse brains directly from the main text, supplementary materials, and/or extrapolated from the figures and graphs presented. In this systematic review, “raw” total immune cell subset counts were reported for each immune cell subset as a median with an interquartile range (IQR; defined as: 75^th^ percentile upper quartile [Q3] – 25^th^ percentile lower quartile [Q1]) and as the combined mean with standard deviation (SD) of multiple studies (n). The “raw” total overall cell count collected per sample by flow cytometry was also reported from these studies.

### 2.5 Data analysis

After the median with IQR and combined means with SD from the “raw” total immune cell subset counts and from the “raw” total overall cell count collected per sample were recorded from each study, we standardized an approach to estimate how many immune cells of each subset would be counted if the total homogenized mouse brain cells collected by flow cytometry equaled a total of 1 x 10^5^ collected cells. The value of cells required, per sample, for an accurate flow cytometry reading is stated to be between 1 x 10^4^ total cells (minimum) and 1 x 10^6^ total cells (maximum) hence we used the median (26).

Standardized total cell counts for each immune cell subset reported in this systematic review, were calculated using the following equation:

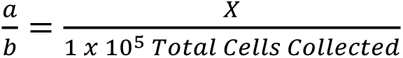

Where “a” is the [Raw Total Immune Cell Subset Count], “b” is the [Raw Total Cell Count Collected] per sample, where “X” is the [Standardized Total Immune Cell Subset Count]. Rearranging to solve for X gives the following equation:

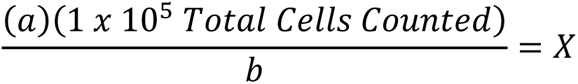

Once all total immune cell subset counts were standardized, we determined the theoretical standardized percentages of each immune cell subset within the entire WT mouse brain, and report those results as means with SD.

## 3 Results

### 3.1 Literature search and study selection

A PRISMA based flow diagram illustrates our screening methodology for study identification (**Figure 1**) (21). The two databases (Google Scholar and PubMed) were searched using keywords, as defined in Methods, and highlighted 900 articles for systematic review. Of those 900 reports, 223 were removed as they supplied only an abstract or were duplicated between databases, and 677 studies remained. Of these an additional 240 studies were removed as they were published in 2012 or prior. This cut off is arbitrary, but was used to identify more contemporary flow cytometry immunophenotyping publications and resulted in 437 eligible studies. Further inclusion and exclusion criteria removed: 133 studies in which flow cytometry of homogenized mouse brains was not a main focus; 129 studies with insufficient information and/or results from controls or in which the background mice strain was not specified; 94 human studies, and; 23 rat studies. Consequently, 58 studies that passed our inclusion and exclusion criteria were incorporated in this systematic review (27–84).

**Figure 1.**
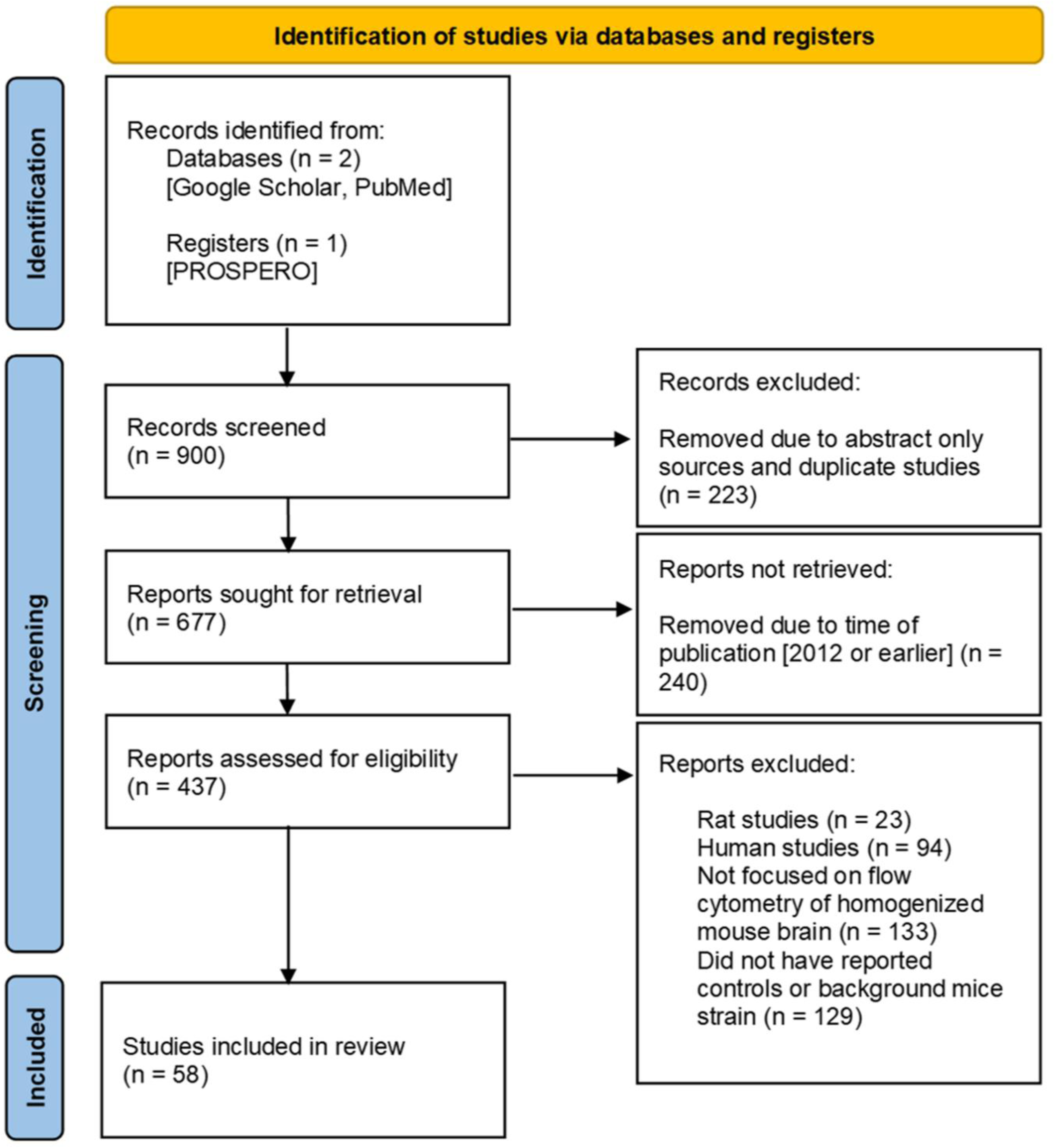
Prisma Flow Diagram. Guidelines provided by PRISMA (21).

### 3.2 Reporting of mouse information and perfusion/tissue processing is inconsistent between studies

Of the 58 studies selected, we reviewed the basic characteristics of the mice strains used (**Figure 2A**, **Suppl. Table 1**). The inbred congenic C57BL/6 mouse line was used most (26/58 studies [44.8%]) as a control and as the background for genetically modified mice (27,28,30,31,36,41–47,49,51,54,58,66–69,72,74–76,79,82). However, other C57BL/6 mice sub-strains were used throughout the 58 studies that include: C57BL/6J (19/58 studies [32.8%]) (29,32,33,39,52,56,60,62–65,70,71,73,78,80,81,83,84); C57/BL6 (2/58 studies [3.44%]) (34,40); C57BL/6 (H-2b) (2/58 studies [3.44%]) (35,53); and C57BL/6J (B6) (2/58 studies [3.44%]) (38,61). The majority of studies that reported mouse sex (**Figure 2B, Suppl. Table 1**) used only male mice (33/58 studies [56.9%]) (29,30,33,34,38,40,41,44,45,47,49,50,54,58–63,65,67–71,74,77–80,82–84), although 10/58 studies [17.2%] reported mixed results of male and female animals together (27,31,32,36,37,39,43,51,72,75). The age of the mice (**Figure 2C, Suppl. Table 1**) within studies varied but the majority reported findings at 8-12 week (2-3 months) (9/58 studies [15.5%]) (29,33,56,57,65,72,75,80,83). However, studies characterized animals over a wide range of age, from 1-2-weeks (32,39,81) and between 3-26 months (37).

**Figure 2.**
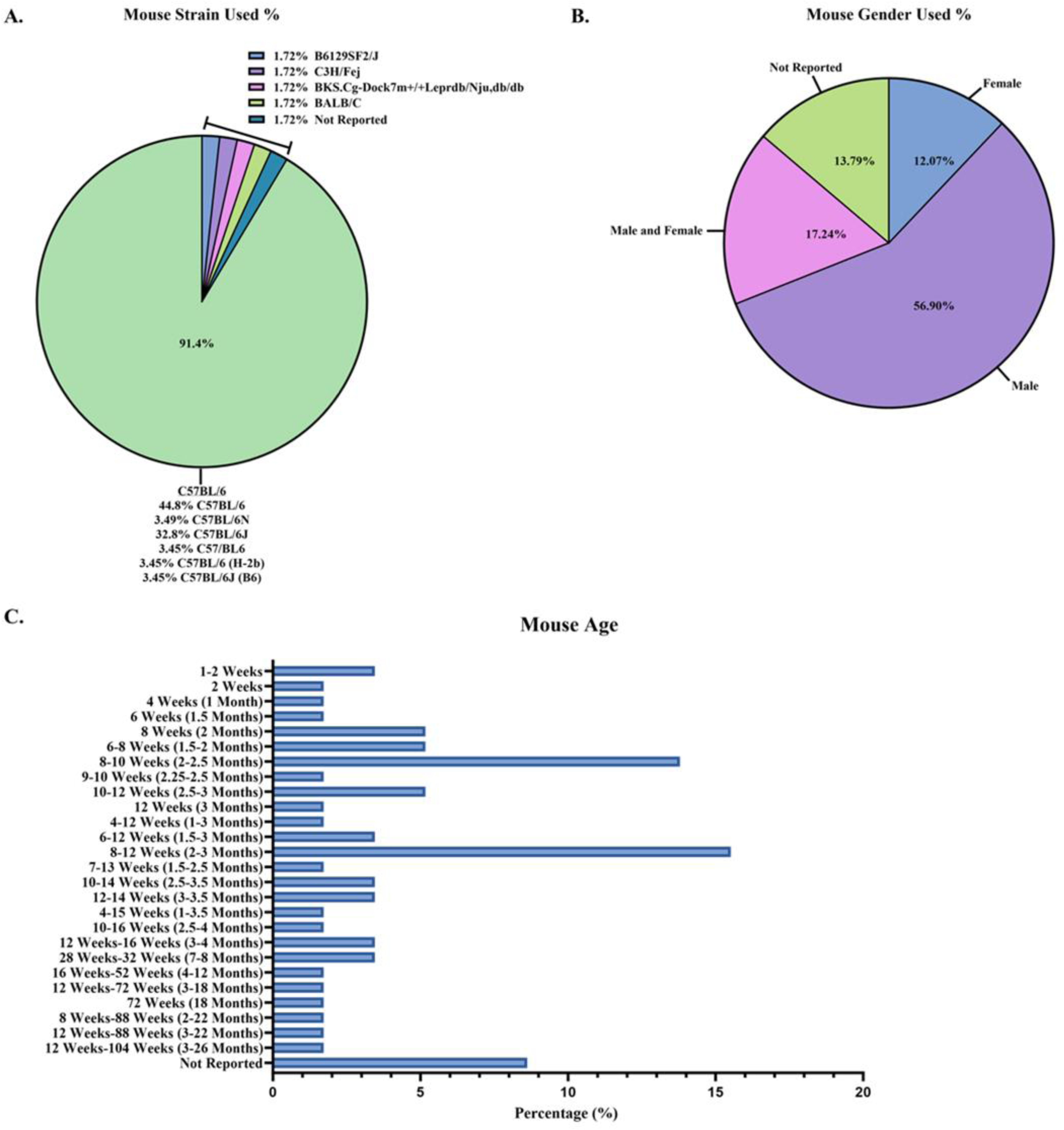
Mouse information reported between studies. Reported baseline mouse information described as percentages out of the 58 studies: **A.** mouse strains; **B.** mouse sex; and **C.** mouse age.

Perfusion (**Suppl. Figure 1, Suppl. Table 1**) and brain tissue processing methods (**Figure 3A-C, Suppl. Table 1**) also varied across the 58 studies. A majority of studies (36/58 studies [62.1%], **Suppl. Figure 1, Suppl. Table 1**) used cold PBS for perfusion (27,29,31–33,35,39,43–48,50–58,61,63–67,70,73–76,79,81,84). Of the 58 studies reviewed, 6/58 [10.3%] (**Figure 3A, Suppl. Table 1**) used a commercial kit, such as the Neural Tissue Dissociation Kit P (Miltenyi Biotec), to isolate immune cells from mouse brain (34,40,41,61,67,71). Nevertheless, in neurology it remains unclear how to best process mouse brain to dissociate the tissue and leave cell types intact (85–87).

**Figure 3.**
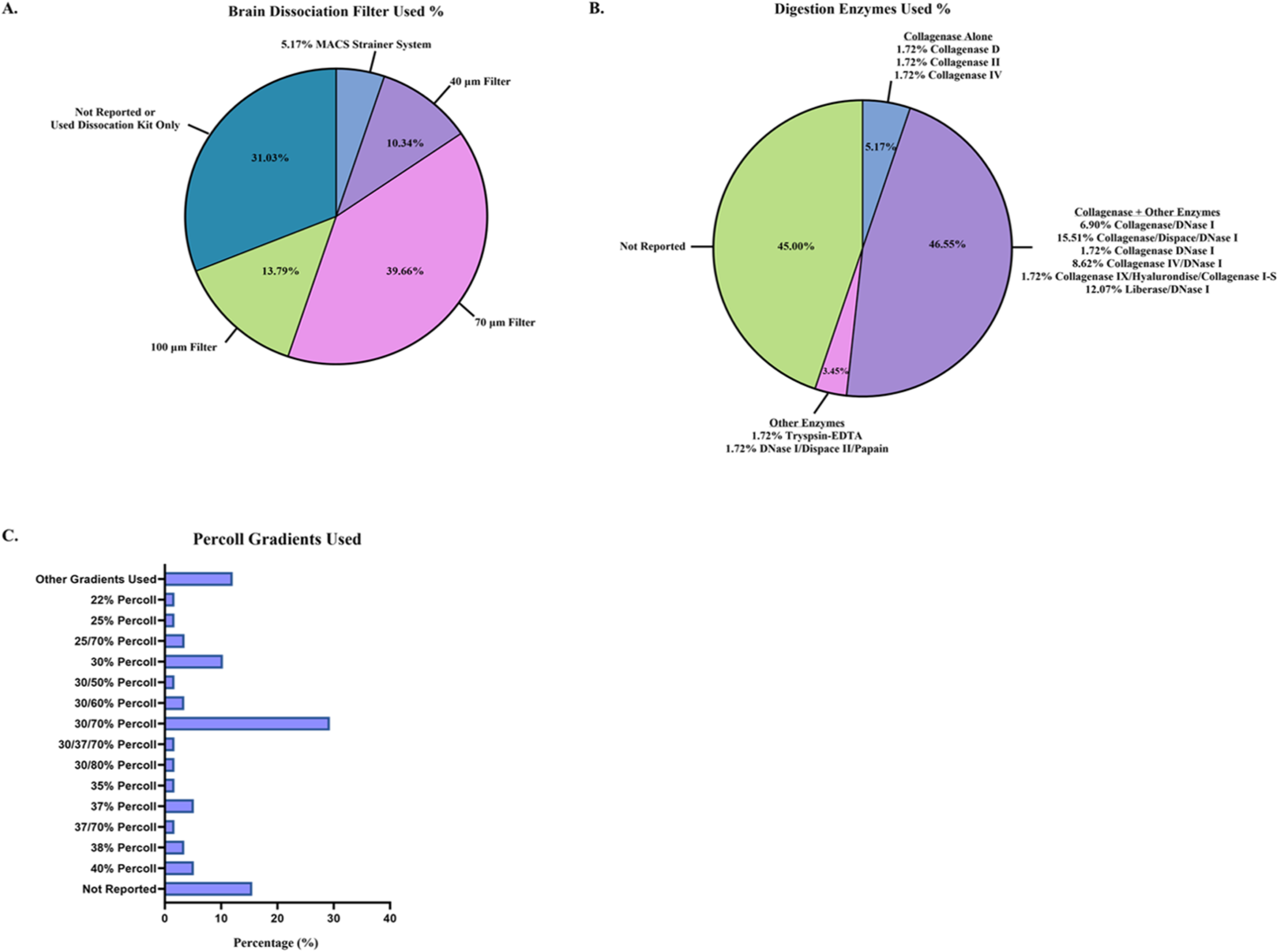
Brain tissue dissociation and cell isolation methods reported between studies. Tissue processing techniques were reported as percentages from the 58 studies: **A.** brain dissociation filters utilized; **B.** digestion enzymes used for brain homogenization; and **C.** Percoll gradients used to remove myelin layer and isolate immune cells.

Researchers may use mechanical homogenization, enzymatic homogenization, or both homogenization techniques (85–87). For the studies reviewed, 18/58 [31.0%] (**Suppl. Table 1**) used some type of mechanical homogenization (i.e. Glass-Teflon homogenizer, an 18-gauge needle, etc.) before filtering through a cell strainer, prior to enzyme treatment and the use of a myelin removal/immune isolation gradient (i.e. Percoll gradient) (31,33,37,44,48,50,53,55–57,59,60,63,64,66,69,81,84). The enzymatic solutions used in the reports also vary widely (**Figure 3B, Suppl. Table 1**). Most studies used collagenase (I, II, IV, D, I-S, or liberase) alone or combined with additional enzymes (30/58 studies [51.7%]) (27–31,33,46–48,53,54,60,62–64,66–68,70,73–80,82–84). The enzyme most used in combination with collagenase (or another tissue digestion enzyme) was DNase I (27/58 studies [46.6%]) (27–31,47,48,51,53,54,60,62,63,66,68,70,73–80,82–84). After homogenization of mouse brain, cell strainers are generally used to remove dead cells and myelin (**Figure 3A, Suppl. Table 1**), and a 70 μm filter was used in the majority (23/58 studies [39.7%]) of studies (15,30,32,33,38,39,43,45,46,48,52,53,56,58,59,62–64,67,72,74,82,84). To further remove myelin from mouse brain homogenate along with isolating immune cells, researchers employ a myelin removal kit or Percoll gradient solutions. Specific cell types can be isolated while the myelin layer rises to the top of the sample tube with centrifuge, depending upon the percentage of Percoll. Again, in the studies reviewed, the percentages of Percoll (**Figure 3C, Suppl. Table 1**) varied with most using a 30%/70% Percoll gradient solution (17/58 studies [29.3%]) (15,27,29,33,43,45,53,57,62,63,71,72,74,77,79,81,84). Overall, the age of mice and methodology for isolating immune cell counts from brain for flow cytometry varied greatly between studies.

### 3.3 Flow cytometry methodology used and cell counts are inconsistently reported between studies

After considering mice strain and brain homogenization methods, we examined the flow cytometry instruments used and data reporting (**Figure 4A-C, Suppl. Figures 2A-B and 3A-B, Suppl. Table 2**). The make and model of flow cytometer/FACS sorter (**Suppl. Figure 2A, Suppl. Table 2**) and analysis software (**Suppl. Figure 2B, Suppl. Table 2**) used in each of studies also varied widely. The flow cytometer most reported was the BD LSRII Flow Cytometer (12/58 studies [20.7%]) (30,32,33,35,36,39,43,51,60,61,63,65) whereas the FACS sorter was the BD FACS Aria III (8/58 studies [13.8%])(31,38,47,49,54,55,64,83). The analysis software most generally used was a version of FlowJo (Tree Star) for flow cytometry immunophenotyping (38/58 studies [65.5%]) (27,29–31,33–36,38,41,43–45,47,49–54,56,58,62,63,65,68–74,76,77,79,81,83,84) and BD FACSDiva specifically for FACS analysis (11/58 studies [19.0%]) (32,35,37,39,42,45,60–62,76,77).

**Figure 4.**
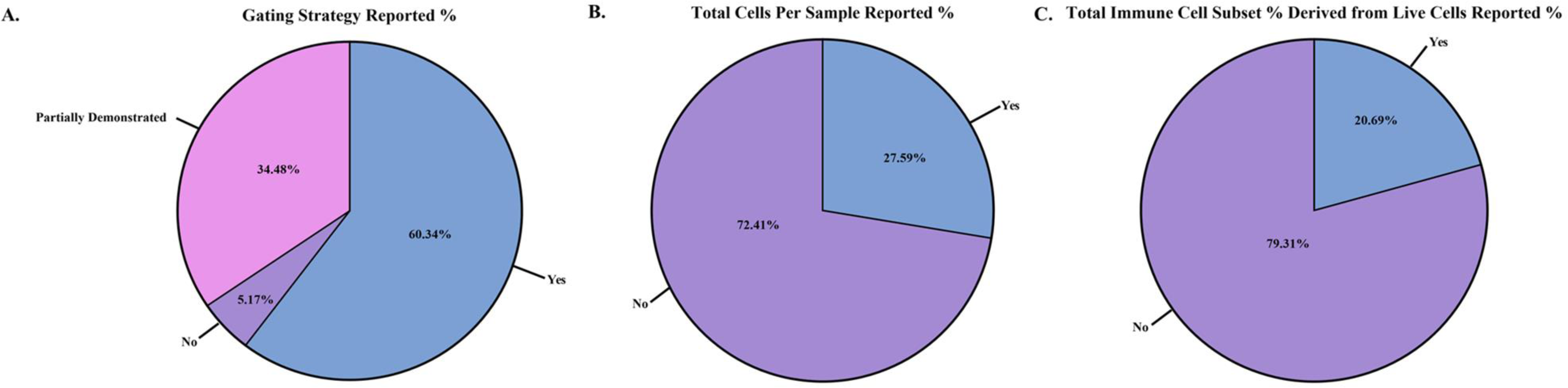
Reporting of flow cytometry results and cell counts between the studies. Differences in flow cytometry results and cell counts reported as within study percentages (n=58 studies): **A.** gating strategy reported; **B.** total cells per sample collected from flow cytometer/sorter reported; and **C.** total immune cell subset counts derived from the live cell gate reported.

After reviewing methodology in all 58 studies, we assessed the quality of flow cytometry reporting (**Figure 4A-C, Suppl. Figure 3A-B, Suppl. Table 2**). For each study, we scored the following parameters: 1) whether a full gating strategy was reported; 2) the total cells collected per sample for flow cytometry; 3) the total live cell counts during flow cytometry sample collection; 4) if the total immune cell subset percentage was calculated directly from the live cell count reported or was it derived from another gate (i.e. was the immune cell subset percentage reported derived from the CD45^+^ gate or from the live/dead gate); and 5) whether the methods used to determine cell counts and/or MFI readings were documented. Of the 58 studies reviewed, 35/58 [60.3%] included a full gating strategy (**Figure 4A, Suppl. Table 2**) (27,30,32–34,36,37,41,43–45,47,50,51,54,55,58,60–64,67,71–79,81,82,84), while 20/58 [34.5%] reported a partial gating strategy (28,29,35,38,40,42,46,48,49,52,53,56,57,59,66,68–70,80,83) and 3/58 [5.17%] of studies did not include this information (31,39,65). When documented, the flow antibody clone and gating strategy was reported (44/58 studies; 75.9%) (**Suppl. Table 2**) (27,29–33,35–40,42,43,45–55,57,58,60–62,66–69,74–80,82–84). Most studies used similar clones to identify specific immune cell subsets (**Supple. Table 2**).

On reporting total cells collected per sample during flow cytometry collection (**Figure 4B, Suppl. Table 2**), only 16/58 [27.6%] of studies provided this data (30,36,37,41,44,47,52,57,58,61,63,67,71,73,74,82). Total live cell counts collected during flow cytometry (**Suppl. Figure 3A, Suppl. Table 2**) was only reported in 9/58 [15.5%] of the studies reviewed (30,37,57,58,61,67,71,73,74). Out of the 58 selected studies, only 12/58 [20.7%] expressed their results in terms of total immune cell subset percentages derived from live cells only (**Figure 4C, Suppl. Table 2**) and not from another gate (such as deriving from the CD45^+^ gate) (35,37,43,47,52,58,67,68,71,72,74,76). Lastly, 35/58 [60.3%] of studies (**Suppl. Figure 3B, Suppl. Table 2**) reported methods on how cell counts and/or MFI readings were recorded (27,28,30,32,33,35,37,39,41–44,49–52,54,55,57–59,61–64,67,70–77,82). Overall, the cytometer/FACS sorter used, and the reporting of total cells collected, total live cells and immune cell subset percentages was not standardized in the 58 studies examined.

### 3.4 Reported immune cell counts from WT/control homogenized mouse brain is highly variable between studies

Across all studies we then examined the total immune cell counts reported in WT/control mouse models, as detected by flow cytometry immunophenotyping and/or by FACS (**Suppl. Table 3**). As described in Methods and **Suppl. Table 3**, studies were selected based on their reporting of a wide-variety of immune cell subsets. These included, T cell subsets/memory T cells (naïve-like, central memory [T_CM_], effector memory [T_EM_] and effector memory re-expressing CD45RA [T_EMRA_]), natural killer cells (NK cells), dendritic cells (DCs), B cells, monocytes, macrophages (M1 [predominately CD86^+^] and M2 [predominately CD163^+^] phenotypes), TMEM119^+^ microglia, and neutrophils. Of note, the M1/M2 nomenclature for macrophages is being updated currently in the immunology field (88)

Many immune subsets were examined but relatively few were reported in sufficient studies to be able to calculate representative medians with IQR and means with SD (**Figure 5 and Suppl. Table 3**). Immune cell subsets with at least two or more references to derive a median and mean cell count for WT/control mice include: bulk CD4^+^ T cells (27–33), bulk CD8^+^ T cells (27,28,30,34–37), double negative (DN) T cells (33,38), regulatory T cells (T_REG_) (29,39–42), T helper 1 cells (T_H1_) (27,40), T helper 17 cells (T_H17_) (44,45), NK cells (33,41,46,47,49–51), DCs (33,52–55), B cells (33,56–60), monocytes (61–67), bulk macrophages (64,65,68–72), TMEM119^+^ microglia (73–76) and neutrophils (46,77–84). Immune cell subset counts that were derived from only one study, such as follicular T cells (T_FH_) (43) and T_EM_ bulk CD4^+^ and CD8^+^ T cells (30), were still included in this review as a representation of the possible median/average cell count for these subsets.

**Figure 5.**
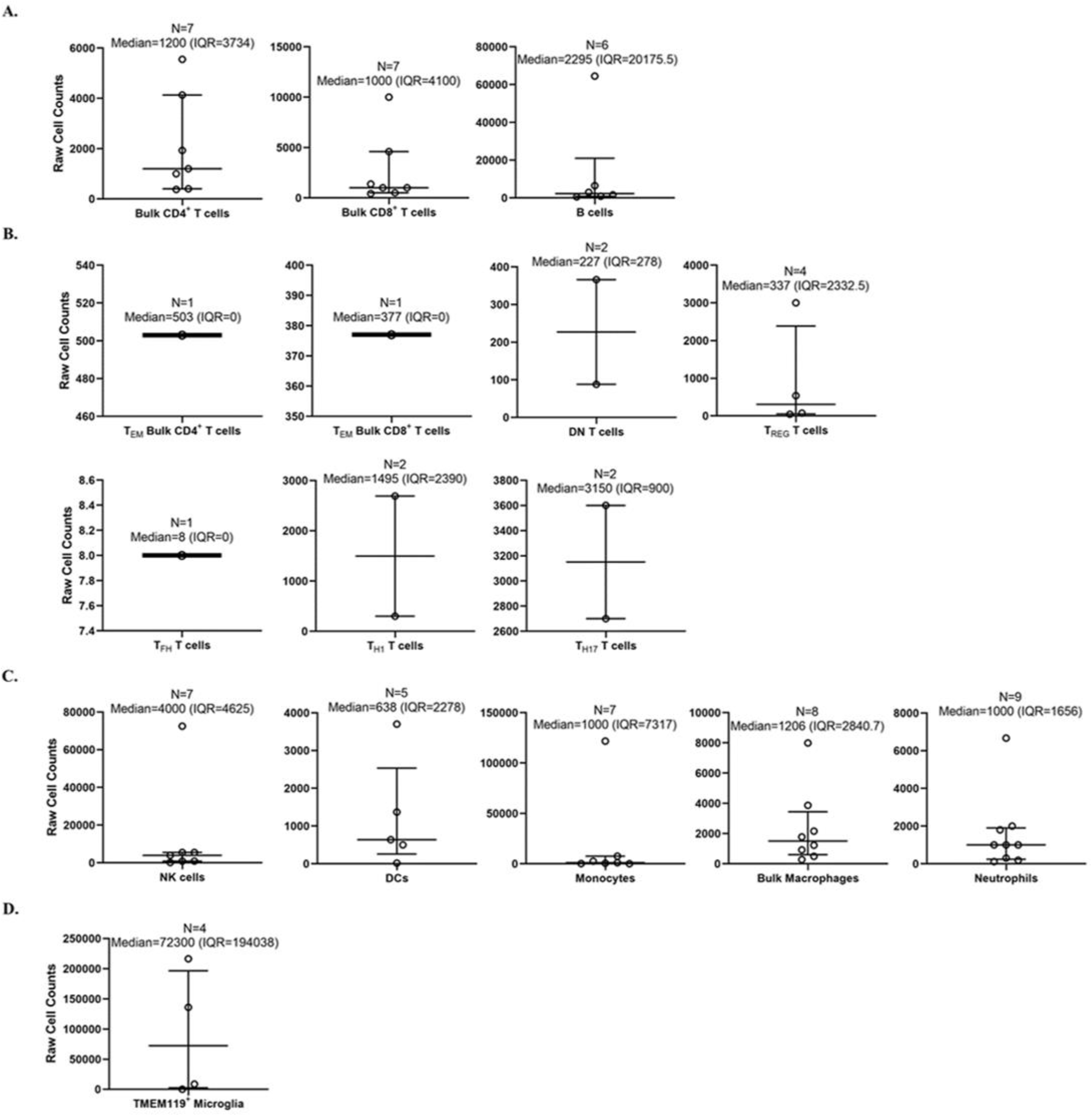
Calculated medians with interquartile ranges (IQRs) of immune cells quantified by flow cytometry within wild-type/control mouse brains. Medians with IQRs (defined as: 75^th^ percentile upper quartile [Q3] – 25^th^ percentile lower quartile [Q1]) of immune cell subset counts found with wild-type (WT)/control homogenized mouse brain were calculated from data extrapolated from the 58 studies selected for inclusion in this systematic review. Immune cell subsets were organized as: **A.** bulk adaptive immune cells; **B.** specialized T cells; **C.** innate immune cells; and **D.** microglia. Total n-values (number of studies for each identified subset) are reported along with the median and IQR for each immune subset above each bar graph.

We first calculated the overall medians for each examined immune subset by collecting all of the “raw” total immune subset cell counts from each of the 58 studies (**Figure 5**). By doing this, we discovered that out of these studies, there were some that were outliers (outside of the IQR) that appear to heavily altered the overall total immune cell counts determined by flow cytometry (27,33,36,41,47,53,58,61,69,72,76,79). Out of all the immune cell subsets from the 58 studies, the highest median was TMEM119^+^ microglia (72,300 [IQR=194,038]; n = 4 studies) (73–76). The lowest median that was calculated was T_FH_ T cells (8 [IQR=0]; n = 1 study) (43). Memory T cell subsets found within both CD4^+^ and CD8^+^ T cells were not reported as an immune cell subset in any study. Similarly, T helper 2 cells (T_H2_) were not reported in any studies.

The 58 studies had highly variable ranges for each immune cell subset found within mouse brain, for which most of the SD calculated were greater than the means (**Suppl. Table 3**). As with the calculated medians, the highest mean cell count reported from the 58 studies was TMEM119^+^ microglia (90,323 ± 104,555; n = 4 studies) (73–76). The lowest mean cell counts reported were for T_FH_ T cells (8 ± 0; n = 1 study) (43). Subsequently, we also calculated the means of total overall cell counts, collected per sample, as reported in the 58 studies reviewed (**Suppl. Table 3**). Although not as variable as immune cell subset counts, the total overall cell counts collected by flow cytometry ranged from 1 x 10^4^ to ∼3 x 10^6^ cells (**Suppl. Table 3**). In some studies, the actual number of cells collected by flow cytometry was not specified, but the total cell counts could be extrapolated from data given in the main text, figures, and/or supplementary material. Overall, we conclude the immune cell subset counts collected by flow cytometry immunophenotyping are highly variable between studies potentially due to the processing method, technical skills and experience of the researcher. Also, there is currently insufficient data on specific T cell subsets/memory subsets and specific macrophage phenotype counts within the WT mouse brain.

### 3.5 Standardizing immune cell counts and percentages within WT/control mouse brain

We devised a method to standardize counts of immune cells and subset percentages within WT/control mouse brain in each study (**Figure 6, Suppl. Table 3**). In flow cytometry, cell type percentages should be reported as values from the total number of cells collected (on average 1 x 10^5^ cells) rather than from a sub-partition within the gating strategy (26,89). Thus, based on the assumption the total cells collected per sample was 1 x 10^5^ cells, we were able to estimate a standardized total immune cell subset count for the data provided in each study.

**Figure 6.**
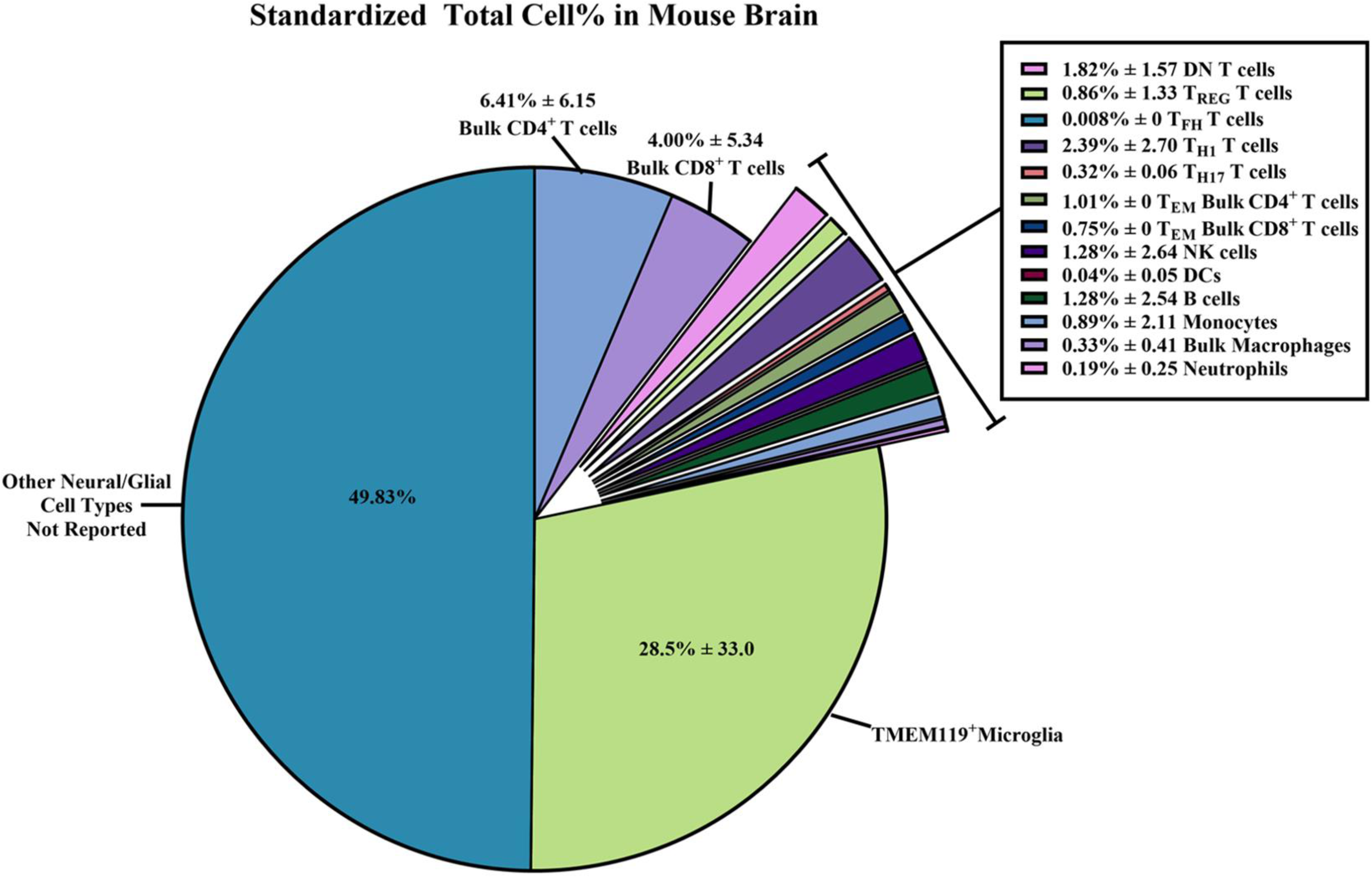
Standardized total cell percentages of immune cells quantified by flow cytometry within wild-type/control mouse brains. Estimated percentages of immune cell subset counts found within wild-type (WT)/control homogenized mouse brain were calculated from data extrapolated from the 58 studies selected for inclusion in this systematic review. The equations used to determine the standardized cell counts can be found in the Methods section. Briefly, “raw” total immune cell subset count and “raw” total cell count collected via flow cytometry were standardized assuming 1 x 10^5^ total cells were collected. Results were reported as combined means of percentages with standard deviations (±SD) from the standardized totals from each study assuming 1 x 10^5^ total cells were collected.

From these standardized total immune cell subset counts, we are able to determine the proportion of immune cell subsets within the brain by simply dividing the standardized count by 1 x 10^5^ total cells collected to obtain percentages (**Figure 6**, **Suppl. Table 3**). The immune subset with the highest total cell percentages that were found within WT/control mouse brain were TMEM119^+^ microglia (28.5% ± 33.0). Of non-neural/glial specific immune cells, bulk CD4^+^ T cells (6.41% ± 6.15) and bulk CD8^+^ T cells (4.00% ± 5.34), were most often counted within mouse brain compared to other adaptive/innate immune cells. Overall, we were able to calculate the average percentage of immune cells found within WT/control mouse brain from the 58 selected studies. Hence, we are able to report a more reliable estimate of the immune cell composition within mouse brain despite their wide SD.

### 3.6 Evaluating risk of bias of all included studies

As per PRISMA and Cochrane criteria for systematic reviews, it is important to evaluate the risk of bias for all studies cited (20,21,24,90). Here we utilize the SYRCLE’s risk of bias tools for animal studies (24) to create a summary graph and “stop-light” figure (**Figure 7**) to highlight the overall bias of each study assessed within the following domains: D1: Sequence Generation (randomization methods used to choose animals for comparable groups); D2: Baseline Characteristics (full description of animal characteristics from all comparable groups); D3: Allocation Concealment (methods used to conceal how animals are distributed to researchers, i.e. using a coding method for each animal); D4: Random Housing (housing all animal groups randomly within the animal room); D5: Blinding (blinding methods used on researchers, such as blinding the knowledge of intervention or transgenic model used and blinding the outcome assessors); D6: Random Outcome Assessment (methods on how the animals were selected at random for outcome assessment); D7: Incomplete Outcome Data (description of the completeness of the data outcome, i.e., stating if data was excluded or if animals were removed from study at any point); D8: Selective Outcome Reporting (the completeness of the study protocols); D9: Other Sources of Bias (examples include: confounders, contamination problems, analysis errors, design-specific risk of bias, etc.).

**Figure 7.**
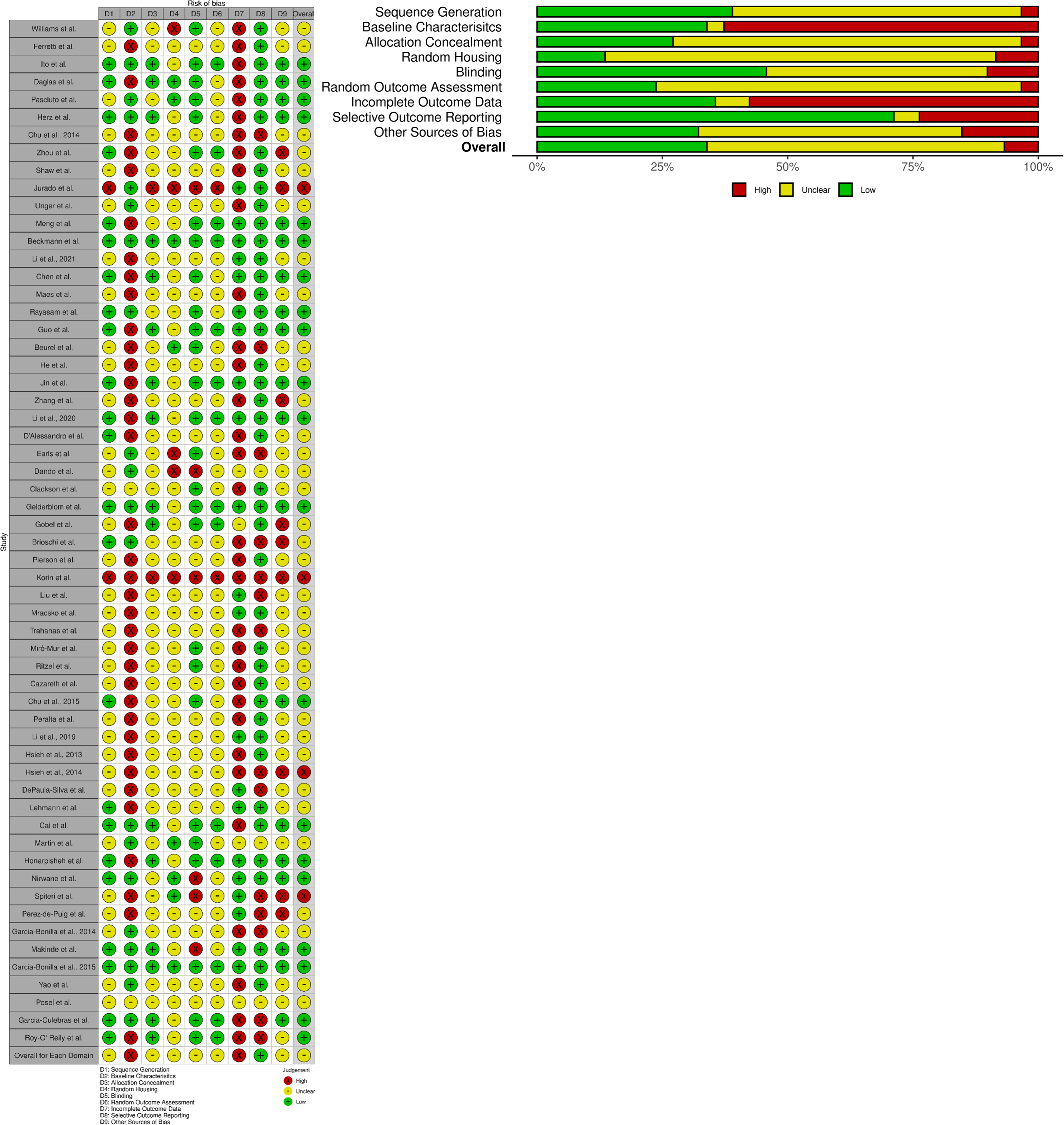
SYRCLE’s risk of bias: Tools for animal studies summarized for each study. Both the summary graph and “stop-light” figure was created via the *robvis* R package and Shiny web app (25). Criteria are based upon the Systematic Review Centre for Laboratory animal Experimentation (SYRCLE) methodology to detect bias (24). All 58 studies selected were reviewed and analyzed using SYRCLE criteria domains: D1: Sequence Generation; D2: Baseline Characteristics; D3: Allocation Concealment; D4: Random Housing; D5: Blinding; D6: Random Outcome Assessment; D7: Incomplete Outcome Data; D8: Selective Outcome Reporting; and D9: Other Sources of Bias. Red: High Bias; Yellow: “Unclear” Bias; Green: Low Bias.

The overall bias of all selected studies was deemed predominantly “unclear” (∼over 50%) due to lack of reporting on specific data/methodology required to pass the “high” or “low” bias questionnaire in each study (**Figure 7**). The most biased domains (∼over 50% high bias scoring) from the selected studies were Baseline Characteristics (i.e. it was largely unclear how sex, age, weight or other baseline characteristics or confounders were adjusted for in each analysis) and Incomplete Outcome Data (i.e. it was generally unclear whether all animals were included in each analysis, and if not, whether there any report on why they were missing outcome data or how that missing data influenced the study). The lowest biased domain (∼over 75% low bias scoring) was Selective Outcome Reporting (i.e. did the results reported reflect on the methods described in the selective studies). For further clarification of methods used in order to clear-up bias reporting, we contacted all 58 corresponding authors to be able to provide us with more information (all names and affiliations of the authors that responded are included in the Acknowledgements section). Overall, the reported bias from all included studies were largely considered to be “unclear” due to the lack of reporting and transparency in the methods and results described. As such, this could be a possible reason for the high variability in immune cell counts that were reported across the multiple studies. Reliable reporting and including confounding factors within experimental procedures/data analysis is necessary for meta-analysis of immunophenotypes found within the mouse brain.

## 4 Discussion and Recommendations

A prominent role for the innate and acquired immune system in brain health, neurologic and age-associated neurodegenerative disorders has become increasing apparent. In part, this has been driven by the immunologic role of several variant gene discoveries including triggering receptor expressed on myeloid cells 2 (*TREM2*) in Alzheimer’s disease, granulin (*GRN*) in frontotemporal dementia and leucine rich repeat kinase 2 (*LRRK2*) in Parkinson’s disease (91–94). Despite this burgeoning interest in neuroimmunology, and many published studies, results from flow cytometry immunophenotyping from homogenized mouse brain are highly variable. Although this does not invalidate ‘within study’ comparisons of specific immune subsets, such variability is a challenge for reproducibility, meta-analysis across studies and interpretation (27–84). Reliable data on residential and infiltrating immune cells within WT/control mouse brain would be of benefit (1–10). One step towards that goal is standardized reporting of flow cytometry methods and results as well as requiring that to become a prerequisite for peer-reviewed publications. In this systematic review, we demonstrate most studies that apply flow cytometry methods to neurology and neuroimmunology, specifically to homogenized mouse brain, share little to no consensus on methods, analysis or results. Here we summarize our findings and make a series of recommendations for future studies (**Table 1**).

**Table 1.**
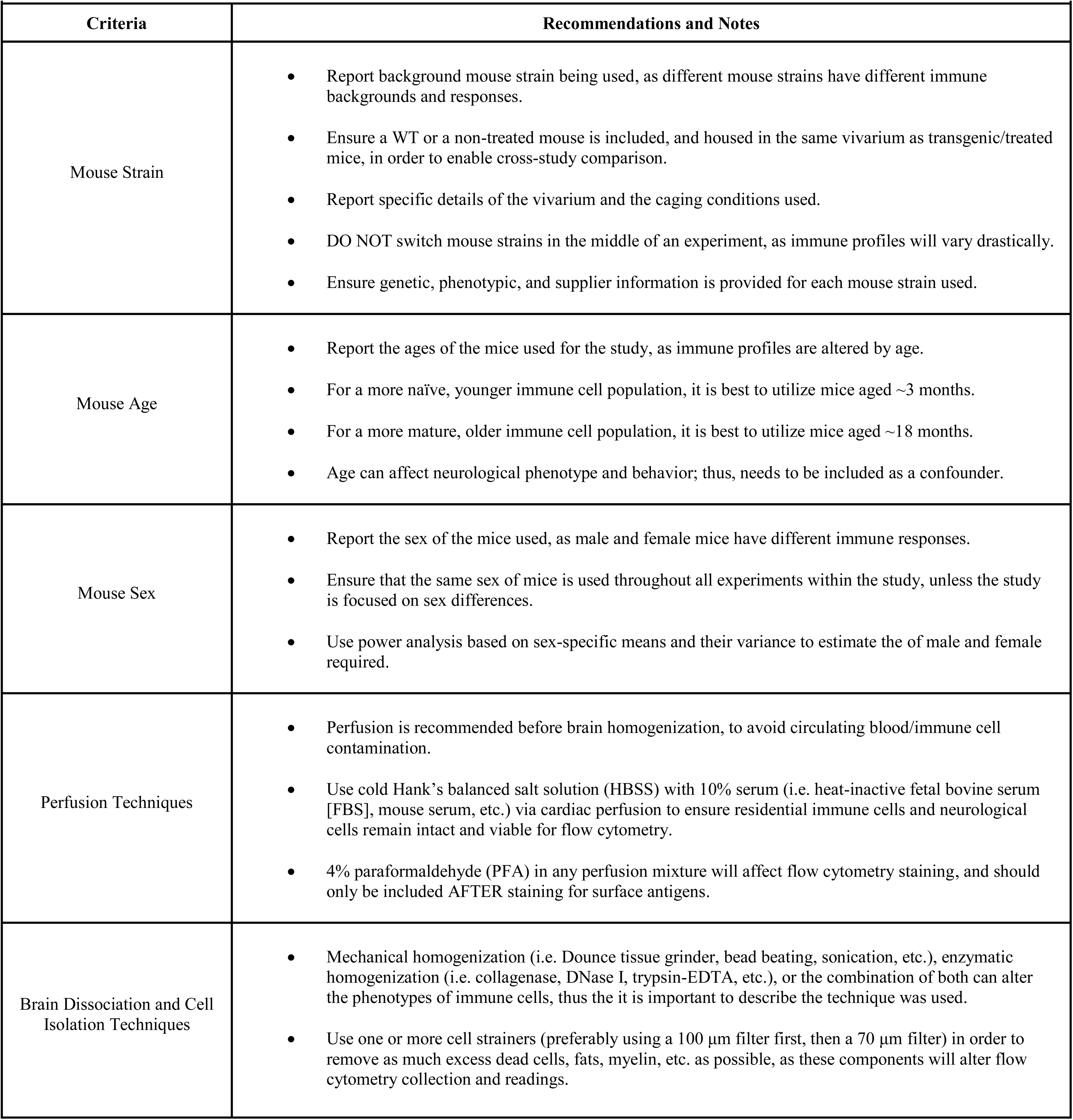
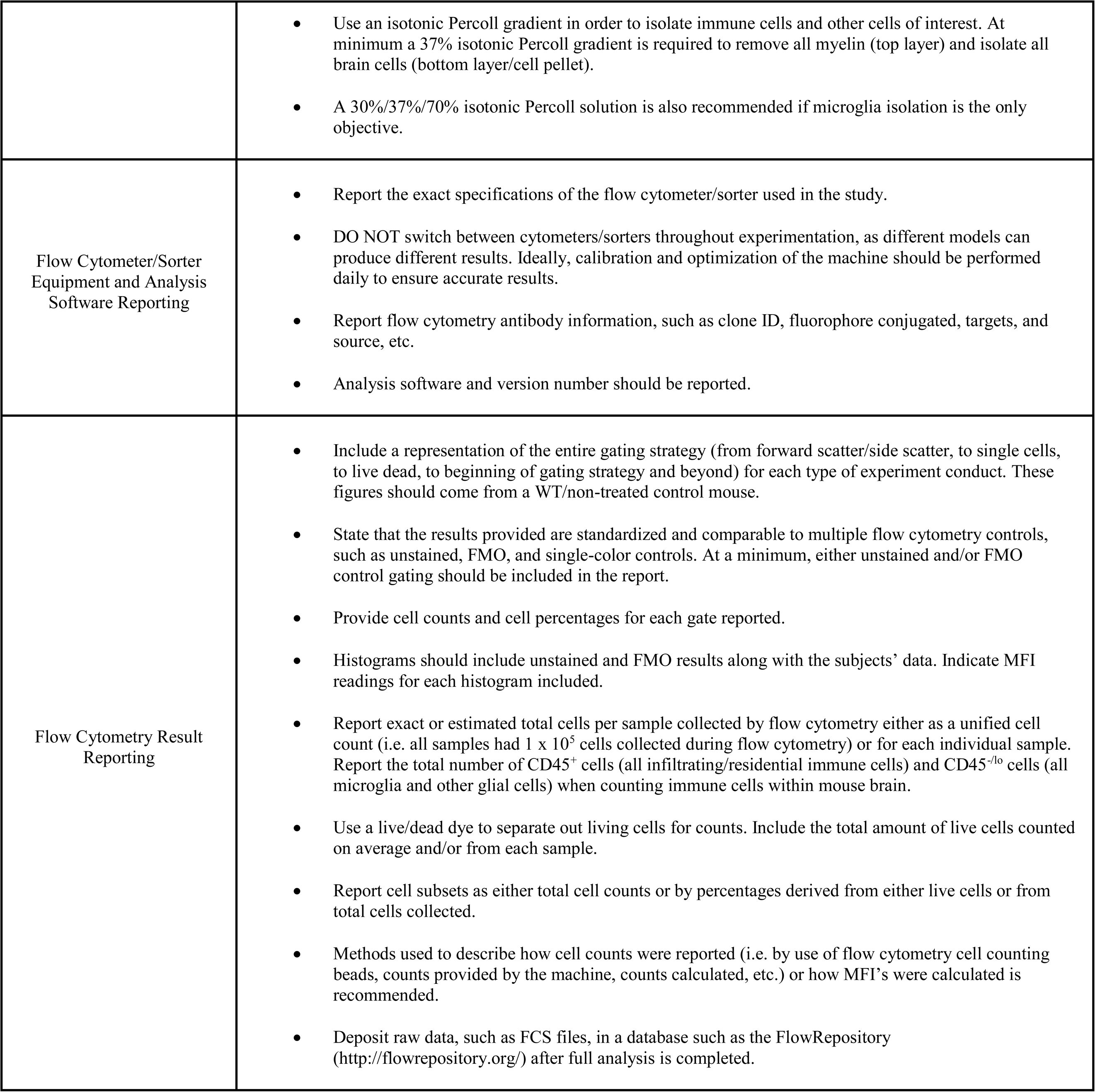
Minimum recommendations and sources for standards for future mouse brain flow cytometry reports (4,5,13,15,26,89,95–98,102,125,126)

We retrieved 58 neurological/neuroimmunology studies that utilized flow cytometry to identify or sort multiple immune cell subsets from WT/control mouse brain, which generally compared within study results to an experimental mouse model (27–84). We compared mouse strains, perfusion and tissue processing methods, and note the age of mice and methods for tissue homogenization are variable (96–99). Vivarium conditions, such as group housing within ventilated racks in a pathogen free barrier facility versus more conventional non-barrier non-ventilated caging, were seldom documented. Corresponding authors from the 10/58 studies (17.2%) indicated the majority of studies utilized a barrier facility with HEPA-filtered air, where each cage was individually ventilated and had sterilized bedding and chow (27,39,45,51,52,58,62,73,75,76). Housing conditions, age and sex influences the immune cell subsets that can be identified by flow cytometry (95–98), and should be carefully considered, documented in experimental protocols and adjusted for as a covariate in subsequent analyses. When deciding on immune cell isolation methods for mouse brains, both mechanical homogenization or/and enzymatic tissue digestion are appropriate. Nevertheless, each approach has pros and cons on immune cell retrieval and phenotypic expression, and depending on the specific research question, must be carefully considered (99–101).

Once the mouse lines and homogenization/isolation methods were analyzed, we compared flow cytometry techniques and data reporting across the 58 studies. Cytometers/FACS sorters come in a variety of makes and models, but essentially do the same function, and should be calibrated using universal standards (89,102,103). Many useful guidelines exist for reporting and include the Minimum Information about a Flow Cytometry Experiment (MIFlowCyt) criteria (26,89,102,104). These recommend researchers present flow cytometry data and methods by reporting: 1) sample/specimen used for experiments; 2) how the samples were treated (storage, processing, stained, etc.); 3) what reagents were used and which antibody clones; 4) what controls were used (unstained controls, FMOs, single color controls, etc.) with a demonstration of the full gating strategy for each panel; 5) what instrument was used and details about it; 6) how many total cells and/or live cells were collected in each sample (either exact cell counts or overall estimated counts); 7) what analysis software was used and how compensation was calculated. Although these guidelines do recommend reporting the total cells and/or live cells collected in each sample, this can be misleading for brain homogenate studies that use different tissue dissociation and cell isolation techniques. For example, a 30%/70% Percoll gradient solution will preferentially isolate immune cells, whereas a 30% solution will isolate immune cells and other residential brain cells (neurons, astrocytes, etc.). Hence, it would be beneficial to report all CD45^+^ (infiltrating/residential immune cells within the brain) and CD45^-/lo^ (microglia and other glial/brain cells). Better documentation would enable replication, more reliable and accurate results, and enable subsequent meta-analysis on immunophenotyping of neurogenerative mouse models.

In the 58 selected studies, we next examined the median and average count and percentages of immune cell subsets reported from WT/control mouse brain. The immune cell subsets selected (**Figure 5**, **Figure 6**, **Suppl. Table 3**) consist of brain/CNS-only residential immune cells (microglia) and immune cells considered to be both residential and peripheral (i.e. T cells, B cells, macrophages, NK cells) (1–10). As expected, of all the immune subsets examined, microglia are the most populous immune cells within WT/control mouse brain (15,74–76). For microglia markers, we searched for publications that showed expression of TMEM119, which is expressed in more stable, non-reactive microglia, (105–108). Traditional methods of detecting microglia by flow cytometry use CD11b^+^ CD45^lo/-^ phenotyping. However, we favor microglia-specific markers, such as TMEM119 as these differentiate microglia from other phagocytic cell types.

For other immune cell subsets, T cells make up the second most abundant immune cell within WT/control mouse brains, with more bulk CD4^+^ T cells (27–37). In the brain, it is likely both CD4^+^/CD8^+^ T cells are comprised of resident memory T cells (T_RM_) and might be classified as an even more specialized subset (i.e. T_CM_, T_EM_, and T_EMRA_) (109–111). For the innate immune cell populations (excluding microglia), NK cells were observed more frequently in WT/control mouse brain than other innate immune cells (33,46–51). Many studies report NK cells as most abundant within the brain’s parenchyma, and more often than other innate immune populations (excluding microglia) or adaptive immune cells (T cell and B cells) (112–114). NK cells were the most reported innate immune cell subset within WT/control mouse brains in the 58 studies reviewed, but there were still more bulk CD4^+^/CD8^+^ T cell counts reported. We cautiously include neutrophils in our systematic review (46,77–84). These polynucleated cells are challenging to detect with flow cytometry as neutrophil extracellular traps (NETs) cause them to be extremely “sticky”, to bind onto each other and to bind non-specifically to flow antibodies (causing false positives) (115–118). The variability in nomenclature/targets to identify neutrophils, their short life span and sensitivity towards purification methods are additional limitations (115–118).

The presence of less abundant immune cell subsets found within the brain, or infiltrating the brain, including specific CD4^+^ T cell subsets, T cell memory subsets, and DCs, were also assessed. Unpredictably, it appears that T_H1_ T cells are more abundant in WT/control mouse brains than T_H2_ T cells (not reported in the 58 selected studies) (27,40). Conventionally, the T_H1_/T_H2_ ratio is used to determine whether an individual has a bacterial/viral infection (higher ratios are indicative or greater infection) although higher ratios of these subsets are also found in aged subjects, once again highlighting the importance of defining age in studies of immunity (119). None of the 58 studies reported specific T cell memory subsets besides bulk T_EM_ CD4^+^/CD8^+^ T cells (30) within WT/control mice. Given the importance of specific T cell memory subsets to overall immunity, future flow cytometry analysis of the brain may benefit from their inclusion. Surprisingly, a few reports listed migratory/residential DCs within WT/control mouse brains, albeit at extremely low levels (33,52–55), as microglia are thought to mediate brain immune surveillance (120–122).

Our systematic review has some limitations. Our analysis and database search were not automated to update figures from more recent research (123,124) and potentially, this may be considered a selection bias. The key word searches we conducted for each immune cell subset has been reported in our methods but different variations of these names, or use of other abbreviations when searching, could alter what literature is identified in each database, thus could also be considered as some level of bias. As several studies selected failed to report sufficient details within their main text, figures, or supplementary materials, whenever possible, data was extrapolated. We have attempted to avoid any author bias or incorrect statements as two independent reviewers assessed all manuscripts. Original authors were also contacted when further clarifications were required.

Bias of each individual study was reported as per guidelines created by multiple organizations that review and conduct systematic reviews (18–24). With a majority of the studies examined, very few were able to clearly state if there was any bias or not within the experimental design. As such, we deemed most of the studies as “unclear” bias due to the lack of actually reporting on specific data/methodology that could pass as “high” or “low” bias. This can be very problematic, as for example, the Baseline Characteristics category of bias was unclear or high in a majority of the 58 studies. Not including mouse baseline characteristics such as sex, age, weight, housing conditions, etc. as confounders of experimentation is extremely problematic, thus can lead to high bias due to the dramatic effect of these factors affecting the immune system of each individual mouse model used. As such, factors like these can heavily affect the results of flow cytometry testing.

We recommend supplementary data includes all raw cell counts and documents which data are used in the main figures and text, to provide more transparency and enable reproducibility in flow cytometry experiments. Overall, there was tremendous variability in immune cell subset counts in the 58 studies reviewed, such that the SD often exceeded mean estimates. Hence medians with IQRs are provided throughout this review. This could be due to a wide variety of reasons, such as the technical skills/experience of the researcher, reporting bias, and confounding factors not considered. There are also methods reporting variability that can contribute towards immune cell count variance across studies such as the following: mouse strain/age/gender, perfusion and brain tissue processing techniques conducted, flow cytometer used, fluorescent antibodies used, and gating strategy utilized. Although heterogeneity in instrumentation and procedures can be unavoidable at some points and is dependent upon the facility where the research is conducted, it would be helpful to standardize some aspects of how immunophenotyping and cell counts are reported. Within study comparisons are still valid, however, researchers should report more robust information about mouse parameters, brain tissue processing, and flow cytometry procedures in order to be replicated in the field. It would be insightful to compare flow cytometry results across studies, not only within them. As such, in our analysis, we have included a series of recommendations to aid interpretation of results, reproducibility and meta-analysis (**Table 1**). Adherence to reporting guidelines will ultimately improve our understanding of the dynamic role of immunity in mouse brain.

## Author Contributions

RCS performed the initial search and analysis of this systematic review, where DTG reviewed all literature to confirm findings and reach a consensus. RCS and MJF contributed equally on the conception, design, organization. All authors contributed to manuscript revision, read, and approved the submitted version.

## Acknowledgments

The authors would like to thank John P.A. Ioannidis from the Meta-Research Innovation Center at Stanford (METRICS) and the Departments of Medicine, Epidemiology and Population Health, Biomedical Data Science, and Statistics at Stanford University, Stanford, CA., U.S.A., for his review and helpful comments. The authors would like to also thank Etty (Tika) Benveniste from the University of Alabama at Birmingham (UAB), Birmingham, AL., U.S.A. and Malu Tansey from the University of Florida, Gainesville, FL., U.S.A., for recommendations on flow cytometry and neuroimmunology. The following corresponding authors from the 58 studies reviewed are thanked for providing additional information about their publications: Ashley S. Harms (University of Alabama at Birmingham, Birmingham, AL., U.S.A.); Josephine Herz (University Hospital Essen, University Duisburg-Essen, Germany); Eleonore Beurel (University of Miami, Miami, FL., U.S.A.); Jae-Kyung Lee (University of Georgia, Athens, GA., U.S.A.); Paul G. McMenamin (Monash University, Clayton, Victoria, Australia); Asya Rolls (Technion-Israel Institute of Technology, Haifa, Israel); Anna M. Planas (Institut d’Investigacions Biomediques August Pi I Sunyer [IDIBAPS] and Institute for Biomedical Research of Barcelona [IIBB]-Spanish Research Council [CSIC], 08036 Barcelona, Spain); Cecile Delarasse (University of Paris, Paris France); Yao Yao (University of South Florida, Tampa, FL., U.S.A.); and Alanna G. Spiteri (University of Sydney, Sydney, Australia). The authors would also like to thank the Farrer lab at the University of Florida for their encouragement and support to perform this systematic review.

## Data Availability Statement

All relevant data is contained within the article: The original contributions presented in the study are included in the article/supplementary material, further inquiries can be directed to the corresponding author/s.

