## Supplemental Figures and Tables for "Experimental procedures for flow cytometry of wild type mouse brain: A systematic review"

**Supplemental Figure 1**

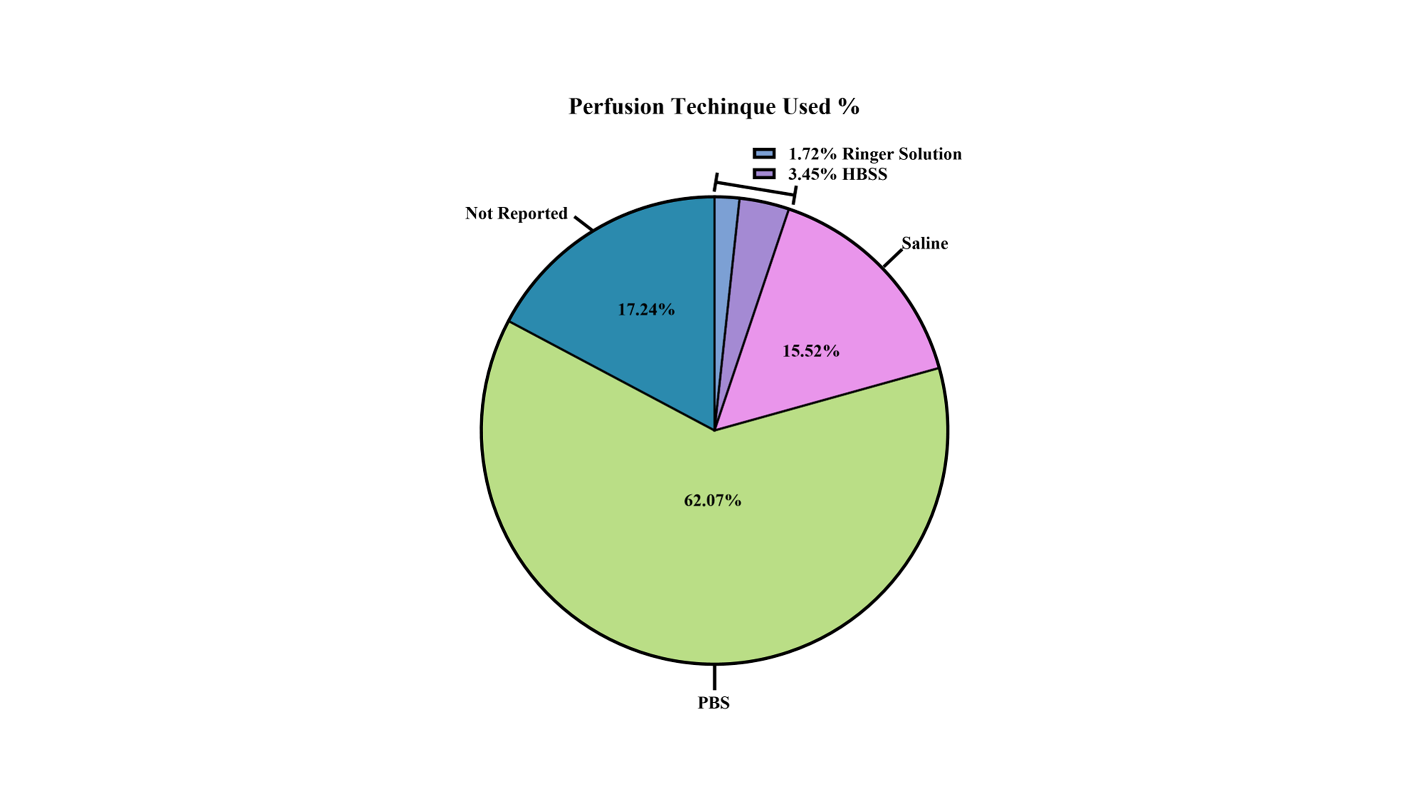

**Supplemental Figure 2**

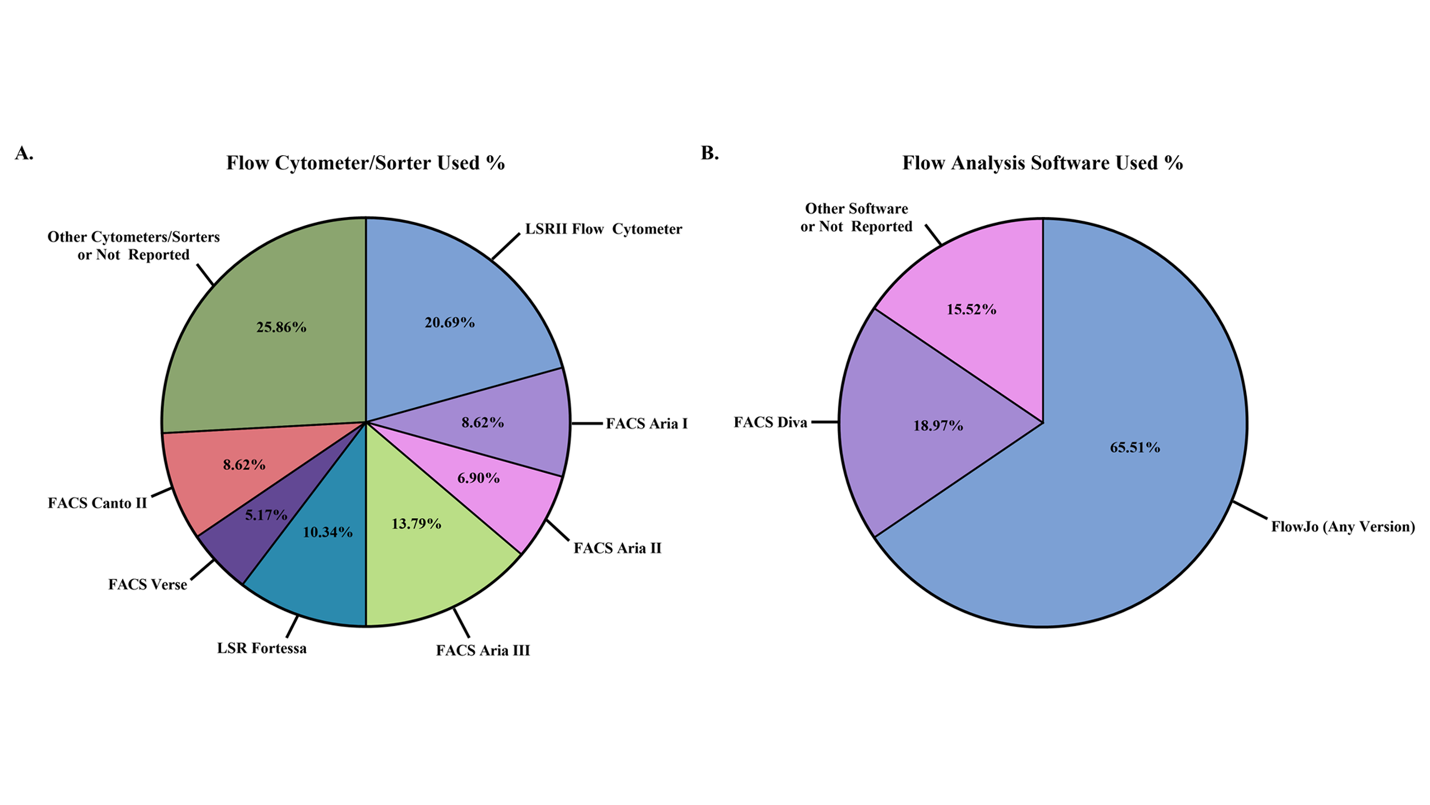

**Supplemental Figure 3**

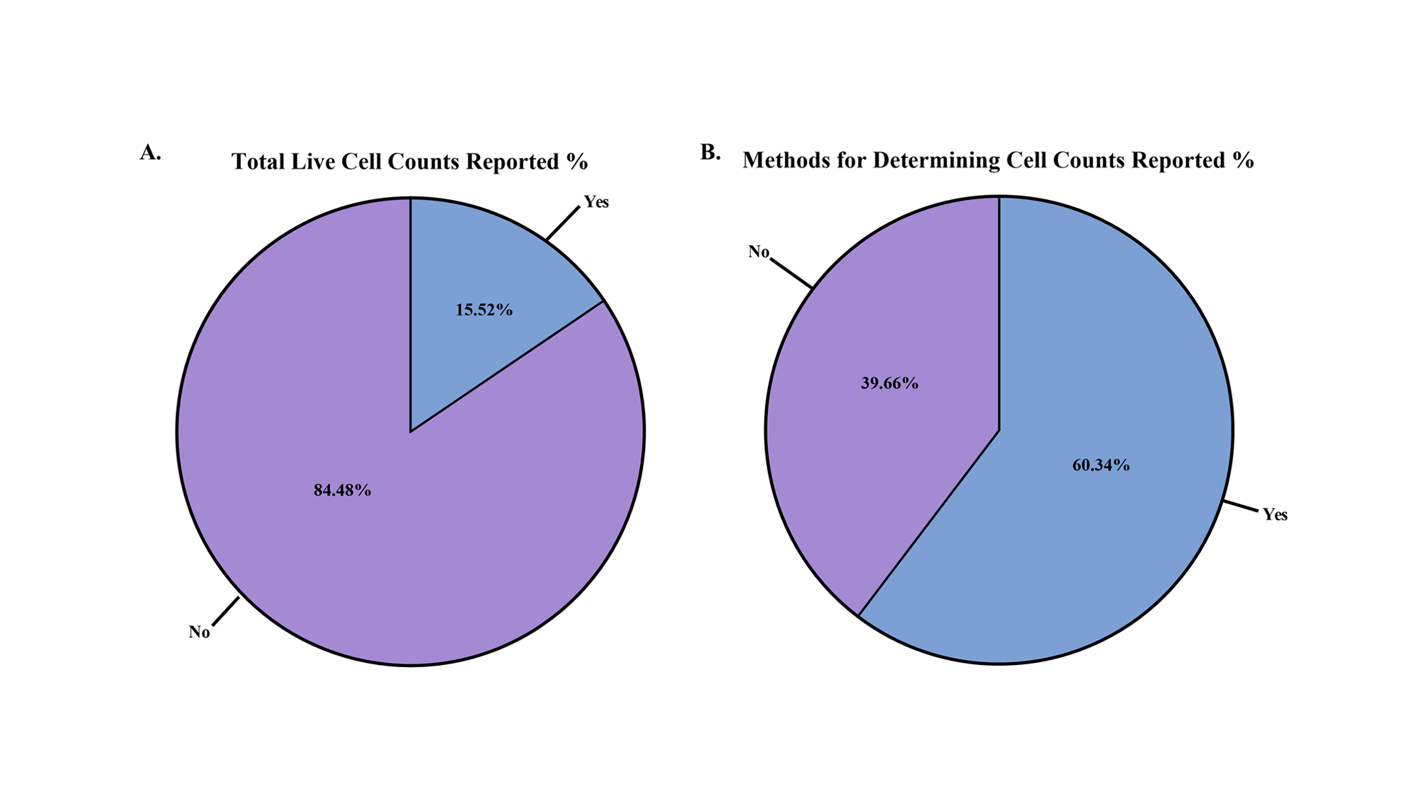

**Supplementary Figure Legends**

**Suppl. Figure 1. Perfusion techniques used on mice for brain tissue processing between the studies.** Perfusion techniques were reported as percentages from the 58 studies.

**Suppl. Figure 2. Flow cytometer/sorter and flow analysis software used between the studies.** Flow cytometers and sorters (**A**) along with flow analysis software (**B**) were reported as percentages from the 58 studies.

**Suppl. Figure 3. Flow cytometry total live cell counts and methods for determining total cell counts used between the studies.** Total live cell counts derived from the live cell gate (**A**) and methods described how cell counts and/or MFI readings were calculated (**B**) were reported as percentages from the 58 studies.

**Supplementary Tables**

**Suppl. Table 1. Wild-type/control mouse information and perfusion/tissue processing reported in references**

| **Suppl. Table 1. Wild-Type/Control Mouse Information and Perfusion/Tissue Processing Reported in References** | | | | | | | |
| --- | --- | --- | --- | --- | --- | --- | --- |
| **Reference** | **Cell Type Examined** | **Mouse Strain WT/Background** | **Age of Mice at Time of Sacrifice** | **Mouse Sex** | **Perfusion Solution** | **Brain Tissue Process Techniques** | **Disease State Studied in Reference** |
| Williams et al. (27) | Bulk CD4^+^ T cells  Bulk CD8^+^ T cells  T_REG_ T cells  T_H1_ T cells | C57BL/6 | 4-12 Weeks | Male and Female | PBS | 1. Collagenase IV (Sigma) Enzyme Solution with DNase I (Sigma)  2. 30/70% Percoll (GE Healthcare) Gradient  3. Filtered Via 100μm Filter | Parkinson’s Disease |
| Ferretti et al. (28) | Bulk CD4^+^ T cells  Bulk CD8^+^ T cells  DN T cells | C57BL/6 | 18 Months | N/R | Ringer Solution (Braun Medical) | 1. DNase I and Collagenase/Dispase (Roche) Enzyme Solution  2. Filtered Via 100μm Filter (Falcon)  3. 30% Percoll (GE Healthcare) Gradient | Alzheimer’s Disease |
| Ito et al. (29) | Bulk CD4^+^ T cells  T_REG_ T cells | C57BL/6J | 8-12 Weeks | Male | PBS | 1. Collagenase D (Roche) and DNase I Enzyme Solution  2. 30/70% Percoll (GE Healthcare) Gradient | Ischaemic Stroke |
| Daglas et al. (30) | Bulk CD4^+^ T cells  Bulk CD8^+^ T cells  T_EM_ Bulk CD4^+^ T cells  T_EM_ Bulk CD8^+^ T cells | C57BL/6 | 7-13 Weeks | Male | N/R | 1. Collagenase/Dispase (Roche), TLCK and Dnase I (Sigma-Aldrich) Enzyme Solution  2. Filtered Via 70μm Filter  3. OptiPrep Solution (Sigma-Aldrich) and MOPS buffer with NaCl Density Gradient | Traumatic Brain Injury |
| Pasciuto et al. (31) | Bulk CD4^+^ T cells | C57BL/6 | 4-15 Weeks | Male and Female | PBS | 1. Mechanical Dissociation  2. Collagenase IV (ThermoFisher), Hyaluronidase (Sigma-Aldrich), and DNase I (Sigma-Aldrich) Enzyme Solution  3. Filtered Via 100μm Filter  4. 40% Percoll (GE Healthcare) | Parkinson’s Disease, Alzheimer’s Disease, and Stroke |
| Herz et al. (32) | Bulk CD4^+^ T cells | C57BL/6J | ~2 Weeks | Male and Female | PBS | 1. Filtered Via 70μm Filter  2. 37% Percoll Gradient | Hypoxic-Ischemic Encephalopathy |
| Chu et al., 2014 (33) | Bulk CD4^+^ T cells  DN T cells  NK cells  DCs  B cells  Bulk Macrophages | C57BL6/J | 8-12 Weeks | Male | PBS | 1. Mechanically Dissociated  2. Collagenase IX, Hyalurondise, and Collagenase I-S in Ca^2+^/Mg^2+^ -Supplemented PBS (Sigma) Enzyme Solution  3. Filtered Via 70μm Filter  4. 30/70% Percoll (GE Healthcare) | Ischaemic Stroke |
| Zhou et al. (34) | Bulk CD8^+^ T cells | C57/BL6 | 8-10 Weeks | Male | Not Directly Reported, Referenced | 1. Neural Tissue Dissociation Kit (Miltenyi Biotec)  2. Percoll Gradient | Ischaemic Stroke |
| Shaw et al. (35) | Bulk CD8^+^ T cells | C57BL/6 (H-2b) | 6-12 Weeks | N/R | PBS | Not Directly Reported, Referenced | Cerebral Malaria |
| Jurado et al. (36) | Bulk CD8^+^ T cells | C57BL/6 | 6-8 Weeks | Male and Female | Not Directly Reported, Referenced | Not Directly Reported, Referenced | Zika Virus Associated Microcephaly and Guillain-Barre Syndrome |
| Unger et al. (37) | Bulk CD8^+^ T cells | N/R | 3 Months-26 Months | Male and Female | HBSS with HEPES (ThermoFisher) and Glucose (Sigma) | 1. Homogenized with Glass Homogenizer  2. Filtered Via 100μm Filter  3. 30% Percoll (Sigma) Gradient | Alzheimer’s Disease |
| Meng et al. (38) | DN T cells | C57BL/6J (B6) | N/R | Male | N/R | 1. “Digestion Buffer”  2. Filtered Via 70μm Filter  3. 37/70% Percoll (GE Healthcare) Gradient | Ischaemic Stroke |
| Beckmann et al. (39) | T_REG_ T cells | C57BL/6J | 1-2 Weeks | Male and Female | PBS | 1. Filtered Via 70μm Filter  2. 37% Percoll Gradient | Neonatal Encephalopathy |
| Li et al., 2021 (40) | T_REG_ T cells  T_H1_ T cells | C57/BL6 | 10 to 12 Weeks | Male | N/R | 1. “Brain Tissue Dissociation Kit” (Miltenyi Biotec) | Parkinson’s Disease |
| Chen et al. (41) | T_REG_ T cells | C57BL/6 | N/R | Male | N/R | 1. Adult Brain Dissociation Kit (Miltenyi Biotec) with gentleMACS Octo Dissociator with Heaters (Miltenyi Biotec) | Ischaemic Stroke |
| Maes et al. (42) | T_REG_ T cells | C57BL/6 | N/R | N/R | Not Directly Reported, Referenced | Not Directly Reported, Referenced | Cancer |
| Rayasam et al. (43) | T_FH_ T cells | C57BL/6 | 10-14 Weeks | Male and Female | PBS | 1. Filtered Via 70μm Filter  2. 30/70% Percoll Gradient | Ischaemic Stroke |
| Guo et al. (44) | T_H17_ T cells | C57BL/6 | 6-8 Weeks | Male | PBS | 1. Mechanically Homogenized and Then Filtered Via 40μm Filter (Becton Dickinson)  2. 30/60% Percoll (GE Healthcare Bio Science AB) Gradient | Ischaemic Stroke |
| Beurel et al. (45) | T_H17_ T cells | C57BL/6 | 6-12 Weeks | Male | PBS | 1. Filtered Via 70μm Filter (BD Biosciences)  2. 30/70% Percoll Gradient | Depression |
| He et al. (46) | NK cells | C57BL/6 | 6-8 Weeks | Female | PBS | 1. Collagenase II (Sigma-Aldrich) Enzyme Solution  2. Filtered Via 70μm Filter  3. 40% Percoll (GE Healthcare) Gradient | Sepsis |
| Jin et al. (47) | NK cells | C57BL/6 | 3-18 Months | Male | PBS | 1. Collagenase IV and DNase Enzyme Solution  2. 30% Percoll Gradient | Aging |
| Zhang et al. (48) | NK cells | B6129SF2/J | 7-8 Months | Female | PBS | 1. Minced with Scissors and Digested with Liberase (Roche) and DNase I (Roche) Enzyme Solution  2. Filtered Via 70μm Filter  3. 40% Percoll (GE Healthcare) Gradient | Alzheimer Disease |
| Li et al., 2020 (49) | NK cells | C57BL/6 | 10-12 Weeks | Male | Not Directly Reported, Referenced | Not Directly Reported, Referenced | Perihematomal Edema |
| D’Alessandro et al. (50) | NK cells | C57BL/6N | 6 Weeks | Male | PBS | 1. Glass-Teflon Homogenizer Passed Through a 100μm Filter  2. 30% Percoll (Sigma) Gradient | Glioma |
| Earls et al. (51) | NK cells | C57BL/6 | 8-10 Weeks | Male and Female | PBS | 1. DNase I (Inivtrogen), Dispace II (Roche), and Papin (Sigma-Aldrich) Enzyme Solution  2. 30/37/70% Percoll (Sigma) Gradient | Parkinson Disease |
| Dando et al. (52) | DCs | C57BL/6J | 8-10 Weeks | N/R | PBS and 1% Heparin | 1. Filtered Via 70μm Filter (Falcon)  2. 30% Percoll (GE Healthcare) Gradient | CNS Regulation |
| Clarkson et al. (53) | DCs | C57BL/6 (H2^b^) | N/R | N/R | PBS and 1% Heparin | 1. Homogenized Through An 18 Gauge Needle  2. Collagenase Type IV and DNase Enzyme Solution  3. Filtered Via 70μm Filter  4. 30/70% Percoll (Pharmacia) Gradient | CNS Autoimmune Diseases |
| Gelderblom et al. (54) | DCs | C57BL/6 | N/R | Male | PBS | 1. Collagenase (Roche) and DNase I (Roche) Enzyme Solution  2. Filtered Via Cell Strainer  3. Percoll (GE Healthcare) Gradient | Ischaemic Stroke |
| Gobel et al. (55) | DCs | C57BL/6N | 8-10 Weeks | Female | PBS | 1. Homogenized  2. 30/50% Percoll (Sigma-Aldrich) Gradient | CNS Autoimmune Diseases |
| Brioschi et al. (56) | B cells | C57BL/6J | 8-12 Weeks;  20-25 Months | Female | PBS | 1. Homogenized  2. Filtered Via 70μm Filter  3. 30% Percoll Gradient | CNS Regulation |
| Pierson et al. (57) | B cells | C3H/Fej | 8-12 Weeks | N/R | PBS | 1. Dissociated Through Sterile Stainless-Steel Mesh  2. 30/70% Percoll Gradient | Multiple Sclerosis |
| Korin et al. (58) | B cells | C57BL/6 | 8-10 Weeks | Male | PBS | Not Directly Reported, Referenced | CNS Regulation |
| Liu et al. (59) | B cells | BKS.Cg-Dock7m+/+ Leprdb/Nju, db/db | 8 Weeks | Male | 0.9% NaCl | 1. Homogenized and Filtered Via 70μm Filter  2. 30/60% Percoll (GE Healthcare) | Ischaemic Stroke |
| Mracsko et al. (60) | B cells | C57BL/6J | 8-10 Weeks | Male | Saline | 1. Mechanically Homogenized  2. Dissociation Buffer (Collagenase and DNase)  3. 1.03/1.088 g/mL Percoll Gradients | Intracerebral Hemorrhage |
| Trahanas et al. (61) | Monocytes | C57BL/6 (B6) | 12-14 Weeks | Male | DPBS | 1. Neural Dissociation Kit (Miltenyi Biotech)  2. Myelin Removal Beads Via AutoMACS Pro (Miltenyi Biotech) | Traumatic Brain Injury |
| Mirό-Mur et al. (62) | Monocytes | C57BL/6J | 3-4 Months | Male | Saline and Heparin | 1. Collagenase IV and DNase I Enzyme Solution  2. Passed Through Tissue Grinder  3. 30/70% Percoll (GE Healthcare) Gradient | Ischaemic Stroke |
| Ritzel et al. (63) | Monocytes | C57BL/6J | 10-12 Weeks | Male | PBS | 1. Mechanically Dissociated and Collagenase, Dispase and DNase (Roche Diagnostics) Enzyme Solution  2. Filtered Via 70μm Filter  3. 30/70% Percoll Gradient | Ischaemic Stroke |
| Cazareth et al. (64) | Monocytes  Bulk Macrophages | C57BL/6J | 8 Weeks | Female | PBS | 1. Homogenized  2. Collagenase D Enzyme Solution (Roche Diagnostics)  3. Filtered Via 70μm Filter (BD Biosciences)  3. 38% Percoll (GE Healthcare) Gradient | Depression |
| Chu et al., 2015 (65) | Monocytes | C57BL/6J | 8-12 Weeks | Male | PBS | Not Directly Reported, Referenced | Ischaemic Stroke |
| Peralta et al. (66) | Monocytes | C57BL/6 | 8 Weeks | Female | PBS | 1. Mechanically Homogenized  2. Collagenase D (Roche Diagnostics) and DNase (Sigma-Aldrich) Enzyme Solution  3. Filtered Via 40μm Filter (BD Biosciences)  4. 38% Percoll (GE Healthcare) Gradient | Parkinson’s Disease |
| Li et al., 2019 (67) | Monocytes | C57BL/6 | 8-10 Weeks | Male | PBS | Dissociation 1  1. Neural Dissociation Kit (Miltenyi Biotech)  2. Myelin Removal Beads  Dissociation 2  1. Collagenase IV (Sigma-Aldrich) Enzyme Solution  2. Filtered Via 70μm Filter  3. Percoll (GE Healthcare) Gradient  4. Filtered Via 70μm Filter | Intracerebral Hemorrhage |
| Hsieh et al., 2013 (68) | Bulk Macrophages | C57BL/6 | 10-16 Weeks | Male | Not Directly Reported, Referenced | 1. Filtered Via 100μm Filter  2. Collagenase Type 1 (Worthington) and DNase I (Sigma-Aldrich) Enzyme Solution  3. Percoll (Amersham Biosciences) Gradient | Traumatic Brain Injury |
| Hsieh et al., 2014 (69) | Bulk Macrophages | C57BL/6 | 10-14 Weeks | Male | Not Directly Reported, Referenced | 1. Mechanically Dissociated and Filtered Via 100μm Filter (BD Biosciences)  2. NOSE Buffer with Collagenase Type I (Worthington) and DNase I (Sigma-Aldrich) Enzyme Solution  3. 1.03/1.095 g/mL Percoll (GE Biosciences) Gradient | Traumatic Brain Injury |
| DePaula-Silva et al. (70) | Bulk Macrophages | C57BL/6J | 4 Weeks | Male | PBS | 1. Collagenase D (Sigma) and DNase I (Roche) Enzyme Solution  2. 37% Percoll (Sigma) Gradient | CNS Regulation |
| Lehmann et al. (71) | Bulk Macrophages | C57BL/6J | 8-10 Weeks | Male | 0.9% Saline | 1. Neural Tissue Dissociation Kit (Miltenyi Biotec)  2. Filtered Via 40μm Filter  3. 30/70% Percoll Gradient | Psychological Stress |
| Cai et al. (72) | Bulk Macrophages | C57BL/6 | 8-12 Weeks | Male and Female | Saline | 1 0.25% Trypsin-EDTA (Thermo Fisher) Enzyme Digestion  2. Filtered Via 70μm Filter (Fisherbrand)  3. 30/70% Percoll (GE Healthcare) Gradient | Ischaemic Stroke |
| Martin et al. (73) | TMEM119^+^ Microglia | C57BL/6J | 4-12 Months | N/R | PBS | 1. “Digestion Cocktail”  2. Filtered Via 70μm Filter  3. 30/70% Percoll Gradient | JoVE Protocol |
| Honarpisheh et al. (74) | TMEM119^+^ Microglia | C57BL/6 | 2-4 Months;  16-22 Months | Male | PBS | 1. Collagenase, Dispase and DNase (Roche Diagnostics) Enzyme Solution  2. Filtered Via 70μm Filter  3. 30/70% Percoll Gradient | CNS Regulation |
| Nirwane et al. (75) | TMEM119^+^ “Microglia | C57BL/6 | 2-3 Months | Male and Female | PBS | 1. 0.2% Collagenase/Dispase (Roche) and DNase I (Sigma) Enzyme Solution  2. Filtered Via 100μm Filter (Fisher Scientific)  3. 22% Percoll (GE Healthcare) Gradient  4. Fixed in 4% PFA | Ischaemic Stroke |
| Spiteri et al. (76) | TMEM119^+^ Microglia | C57BL/6 | 9-10 Weeks | Female | PBS | 1. Collagenase (Sigma-Aldrich) and DNase I (DN25) Enzyme Solution  2. GentleMACS Dissociator (Miltenyi Biotec)  3. 30/80% Percoll Gradient | West Nile Virus Encephalitis |
| Perez-de-Puig et al. (77) | Neutrophils | BALB/C | 3-4 Months | Male | Saline | 1. Collagenase IV and DNase I Enzyme Solution  2. Tissue Grinder  3. 30/70% Percoll Gradient (GE Healthcare) | Ischemic Stroke |
| Garcia-Bonilla et al., 2014 (78) | Neutrophils | C57BL/6J | 7-8 Weeks | Male | Saline | 1. Gentle MACS Dissociator (Miltenyi Biotec)  2. Liberase DH (Roche Diagnostics) and DNase I Enzyme Solution  3. 25/70% Percoll (GE Healthcare) Gradient | Ischemic Brain Injury |
| Makinde et al. (79) | Neutrophils | C57BL/6 | 12-14 Weeks | Male | PBS | 1. Liberase TL (Roche) and DNase I Enzyme Solution  2. MACS Dissociator  3. Filtered Via 40μm Filter  4. 30/70% Percoll Gradient | Traumatic Brain Injury |
| Garcia-Bonilla et al., 2015 (80) | Neutrophils | C57BL/6J | 8-12 Weeks | Male | Saline | 1. Gentle MACS Dissociator (Miltenyi Biotec)  2. Liberase DH (Roche Diagnostics) and DNase I Enzyme Solution  3. 25/70% Percoll (GE Healthcare) Gradient | Ischaemic Stroke |
| Yao et al. (81) | Neutrophils | C57BL/6J | 10-Day-Old | N/R | PBS | 1. Gently Homogenized  2. Filtered via 40μm Filter  3. 30/70% Percoll (GE Healthcare) Gradient | Hypoxic-Ischemic Brain Injury |
| Posel et al. (82) | Neutrophils | C57BL/6 | 12 Weeks | Male | HBSS | 1. Filtered Via 100μm Filter  2. Liberase TL and DNase I Enzyme Solution  3. Filtered Via 70μm Filter  4. 25% Percoll Gradient | JoVE Protocol |
| Garcia-Culebras et al. (83) | Neutrophils | C57BL/6J | 8-12 Weeks | Male | Saline | 1. Collagenase, Dispase, Na-Tosyl-L-Lysine Chloromethyl Ketone Hydrochloride, and DNase I Enzyme Solution  2. Homogenization With Dounce Homogenizer  3. 35% Percoll (GE Healthcare) Gradient | Stroke |
| Roy-O’Reily et al. (84) | Neutrophils | C57BL/6J | 3-4 Months; 20-22 Months | Male | PBS | 1. Collagenase/Dispase (Roche Diagnostics), and DNase (Roche Diagnostics) Enzyme Solution  2. Filtered Via 70μm Filter  3. 30/70% Percoll (GE Lifesciences) Gradient | Ischemic Stroke |
| N/R: Not Reported | | | | | | | |

**Suppl. Table 2. Wild-type/control mouse brain immune cell flow cytometry techniques reported for references**

|  | **Suppl. Table 2. Wild-Type/Control Mouse Brain Immune Cell Flow Cytometry Techniques Reported for References** | | | | | | | | |
| --- | --- | --- | --- | --- | --- | --- | --- | --- | --- |
| **Reference** | **Cell Type Examined for This Study** | **Cytometer(s) Used** | **Software Used** | **Full Gating Strategy?** | **Flow Antibody Clones Used for Targeting Cell Types** | **Total Cells Collected Per Sample Reported?** | **Total Live Cell Counts Reported?** | **Total Immune Cell Subset % Calculated Directly from Live Cell Count Reported?** | **Methods Reported for Determining Cell Counts and/or MFI Readings?** |
| Williams et al. (27) | Bulk CD4^+^ T cells  Bulk CD8^+^ T cells  T_REG_ T cells  T_H1_ T cells | Attune Nxt (Thermo Fisher Scientific) | FlowJo (Tree Star) | Yes | CD4^+^ T cells: GK1.5, BioLegend  CD8^+^ T cells: 53-6.7, BioLegend  T_REG_ T cells (FOXP3): FJK-16s, eBioscience  T_H1_ T cells (T-bet): 4B10, BioLegend | No | No | No | Yes |
| Ferretti et al. (28) | Bulk CD4^+^ T cells  Bulk CD8^+^ T cells  DN T cells | BD LSR Fortessa | N/R | P/D | N/R | No | No | No | Yes |
| Ito et al. (29) | Bulk CD4^+^ T cells  T_REG_ T cells | FACSAria IIu (Becton Dickinson);  FACSCanto II (Becton Dickinson) | FlowJo (Tree Star) | P/D | CD4^+^ T cells: RM4-5, BioLegend or eBioscience  T_REG_ T cells (FOXP3): FJK-16s, eBioscience | No | No | No | No |
| Daglas et al. (30) | Bulk CD4^+^ T cells  Bulk CD8^+^ T cells  T_EM_ Bulk CD4^+^ T cells  T_EM_ Bulk CD8^+^ T cells | BD LSRFortessa;  LSRII Flow Cytometer (BD Biosciences) | FlowJo VX (Tree Star) | Yes | CD3^+^ T cells: 17A2, eBioscience  CD4^+^ T cells: RM4-5, BD Biosciences; RM4-4, BioLegend  CD8^+^ T cells: 53-6.7, eBioscience; 53-5.8, Biolegend  T_EM_ T cells (CD62L): MEL-14, eBioscience  T_EM_ T cells (CD44): IM7, eBioscience | Yes | Yes | No | Yes |
| Pasciuto et al. (31) | Bulk CD4^+^ T cells | BD FACSymphony;  BioRad ZE5;  BD FACSAria III | FlowJo and R (version 3.6.2) | No | CD4^+^ T cells: GK1.5, BioXCell, BD Biosciences; RM4-4, BioLegend, BD Biosciences | No | No | No | No |
| Herz et al. (32) | Bulk CD4^+^ T cells | BD FACS LSRII | FACS Diva (BD Biosciences) | Yes | CD4^+^ T cells: RM4-5, BD Biosciences | Not Directly, Referenced | Not Directly, Referenced | No | Yes |
| Chu et al., 2014 (33) | Bulk CD4^+^ T cells  DN T cells  NK cells  DCs  B cells  Bulk Macrophages | LSRII (BD Biosciences) | FlowJo (Tree Star) | Yes | CD3^+^ T cells: 17A2, eBioscience  CD4^+^ T cells: GK1.5, BioLegend  NK cells (NK1.1): PK136, BioLegend  DCs: N418, BioLegend  B cells (B220): RA3-6B2, BioLegend  Macrophages (F4/80): BM8, eBioscience | No | No | No | Yes |
| Zhou et al. (34) | Bulk CD8^+^ T cells | FACS Verse (BD Biosciences) | FlowJo (Tree Star) | Yes | N/R | No | No | No | No |
| Shaw et al. (35) | Bulk CD8^+^ T cells | BD LSR II (Becton Dickinson);  MACSQaunt (Miltenyi) | BD FACSDiva (Becton Dickinson) and FlowJo (Tree Star) | P/D | CD8^+^ T cells: 53-6.7, N/R | Not Directly, Referenced | Not Directly, Referenced | Yes | Yes |
| Jurado et al. (36) | Bulk CD8^+^ T cells | LSR II (BD Biosciences) | FlowJo (Tree Star) | Yes | CD8^+^ T cells: 53-6.7, eBioscience or BioLegend | Yes | No | No | No |
| Unger et al. (37) | Bulk CD8^+^ T cells | LSR Fortessa (BD) | BD FACSDiva (8.0.1, BD);  Kaluza (1.3; Beckman Coulter) | Yes | CD8^+^ T cells: H53-17.2, eBioscience | Yes | Yes | Yes | Yes |
| Meng et al. (38) | DN T cells | Accui C6 (BD Biosciences);  BD FACSAria III (BD Biosciences) | FlowJo v 10 (Tree Star) | P/D | DN T cells (CD3): 145-2C11, eBioscience  DN T cells (CD4): GKI 1.5, eBioscience  DN T cells (CD8): 53-6.7 eBioscience | No | No | No | No |
| Beckmann et al. (39) | T_REG_ T cells | FACS Aria II (BD Biosciences);  LSRII (BD Biosciences) | BD Diva (BD Biosciences) | No | T_REG_ T cells (CD3): 17A2, eBioscience  T_REG_ T cells (CD4): RM4-5, BD Biosciences  T_REG_ T cells (FOXP3): FJK-16s, eBioscience | No | No | No | Yes |
| Li et al., 2021 (40) | T_REG_ T cells  T_H1_ T cells | Attune NxT (Applied Biosystems/Thermo Fisher) | Kaluza 2.0 (Beckman Coulter) | P/D | T_REG_ T cells and T_H1_ T cells (CD3): 145-2C1, eBioscience  T_REG_ T cells and T_H1_ T cells (CD4): RM4-5, eBioscience  T_REG_ T cells (CD25): PC61.5, eBioscience  T_REG_ T cells (FOXP3): FJK-16s, eBioscience  T_H1_ T cells (T-bet): 4B10, eBioscience | No | No | No | No |
| Chen et al. (41) | T_REG_ T cells | BD C6 (Becton Dickinson) | FlowJo | Yes | N/R | Yes | No | No | Yes |
| Maes et al. (42) | T_REG_ T cells | FacsCanto II (BD Biosciences) | FacsDiva (BD) | P/D | T_REG_ T cells (CD4): GK 1.5, eBioscience  T_REG_ T cells (CD25): 7D4, eBioscience  T_REG_ T cells (FOXP3): cFJK-16s, eBioscience | No | No | No | Yes |
| Rayasam et al. (43) | T_FH_ T cells | BD LSR II (BD Biosciences) | FlowJo (Tree Star) | Yes | T_FH_ T cells (CD4): RM4-5, BD Biosciences  T_FH_ T cells (ICOS-1): D10.G4.1, BD Biosciences  T_FH_ T cells (CXCR5): 2G8, BD Biosciences  T_FH_ T cells (IL-21): 06-1074, EDM Millipore | No | No | Yes | Yes |
| Guo et al. (44) | T_H17_ T cells | FACS Aria (BD Biosciences) | FlowJo 7.6.1 (Tree Star) | Yes | N/R | Yes | No | No | Yes |
| Beurel et al. (45) | T_H17_ T cells | FACS Diva; FlowJo | N/R | Yes | T_H17_ T cells (CD4): GK1.5, eBioscience  T_H17_ T cells (IL-17A): eBio17B7, eBioscience | No | No | No | No |
| He et al. (46) | NK cells | BD FACS Aria II | N/R | P/D | NK cells (CD45): 30-F11, eBioscience  NK cells (NK1.1): PK136, eBioscience | No | No | No | No |
| Jin et al. (47) | NK cells | FACSAria III | FlowJo | Yes | NK cells (CD45): 30-F11, BD Biosciences  NK cells (NK1.1): PK136, BD Biosciences | Yes | No | Yes | No |
| Zhang et al. (48) | NK cells | FACSAria II (BD Biosciences) | FACSCanto (BD Biosciences) | P/D | NK cells (CD45): 104, BioLegend or Thermo  NK cells (NK1.1): PK136, BioLegend or Thermo | No | No | No | No |
| Li et al., 2020 (49) | NK cells | FACS Aria III (BD Bioscience) | FlowJo v7.6 (Informer Technologies) | P/D | NK cells (CD45): 30-F11  NK cells (NK1.1): PK136 | No | No | No | Yes |
| D’Alessandro et al. (50) | NK cells | FACS-Cantoll (BD Biosciences) | FlowJo 9.3.2 (Tree Star) | Yes | NK cells (CD45): 104, eBioscience  NK cells (NK1.1): PK136, eBioscience | No | No | No | Yes |
| Earls et al. (51) | NK cells | LSR II (BD Biosciences) | FlowJo 10.0.8 | Yes | NK cells (NK1.1): PK136 | No | No | No | Yes |
| Dando et al. (52) | DCs | LSR Fortessa X-20 (BD Biosciences) | FlowJo 10.1 (Tree Star) | P/D | DCs: HL3, BD Biosciences | Yes | No | Yes | Yes |
| Clarkson et al. (53) | DCs | FACSCalibur;  LSRII (BD Biosciences) | FlowJo 10.0.6 (Tree Star) | P/D | DCs (CD45): 30-F11, BD Biosciences  DCs (CD11c): HL3, BD Biosciences | No | No | No | No |
| Gelderblom et al. (54) | DCs | Fortessa FACS (BD Biosciences);  BD FACS Aria IIIu | FlowJo (Tree Star) | Yes | DCs (CD45): 30-F11, eBioscience  DCs (CD11b): M170, eBioscience  DCs (CD11c): N418, eBioscience | Not Directly, Referenced | No | No | Yes |
| Gobel et al. (55) | DCs | BD FACS Aria III;  FACSGallios | Kaluza (Beckman Coulter) | Yes | DCs (CD11c): N418, BioLegend | No | No | No | Yes |
| Brioschi et al. (56) | B cells | BD X20;  BD LSR Fortessa;  BD Canto-II  (BD Bioscience) | FlowJo 10 | P/D | B cells (CD45): 30-F11, BioLegend  B cells (CD19): 6D5, BioLegend | No | No | No | No |
| Pierson et al. (57) | B cells | FACSAria (BD Biosciences) | N/R | P/D | B cells (CD45): 30-F11, eBioscience  B cells (MHC-II):11-5.2, BD Biosciences  B cells (CD19): 1D3, eBiosciences | Yes | Yes | No | Yes |
| Korin et al. (58) | B cells | LSRFortessa (BD Biosciences) | FlowJo 10.1r5 (Tree Star);  Cytobank | Yes | B cells (CD45): 30-F11, BioLegend  B cells (CD19): 6D5, BioLegend | Yes | Yes | Yes | Yes |
| Liu et al. (59) | B cells | FACSCalibur (Becton Dickinson) | CellQuest (Becton Dickinson) | P/D | N/R | No | No | No | Yes |
| Mracsko et al. (60) | B cells | LSR II (Becton Dickinson) | Diva | Yes | B cells (CD45): A20 or 104  B cells (B220): RA3-6B2 | No | No | No | No |
| Trahanas et al. (61) | Monocytes | LSR II (BD Immunocytometry Systems) | FACS Diva (BD Biosciences) | Yes | Monocytes (CD45): 30-F11, BD Biosciences  Monocytes (CD11b): M1-70, eBioscience  Monocytes (CD64): x54-5/7.1, BioLegend | Yes | Yes | No | Yes |
| Mirό-Mur et al. (62) | Monocytes | FACSCanto II (BD Biosciences) | FACSDiva (BD Biosciences);  FlowJo 7.6.5 (TreeStar) | Yes | Monocytes (CD11b): M1/70 BD Pharmingen  Monocytes (Ly6C): ER-MP20, eBioscience  Monocytes (CD43): S7, BD Pharmingen | No | No | No | Yes |
| Ritzel et al. (63) | Monocytes | LSRII (BD Biosciences) | FlowJo (Tree Star) | Yes | N/R | Yes | No | No | Yes |
| Cazareth et al. (64) | Monocytes  Bulk Macrophages | FACS Aria III (BD Biosciences) | N/R | Yes | N/R | No | No | No | Yes |
| Chu et al., 2015 (65) | Monocytes | LSRII (BD Biosciences) | FlowJo (Tree Star) | No | N/R | No | No | No | No |
| Peralta et al. (66) | Monocytes | FACSCanto II (BD Biosciences) | FCS Express (De Novo) | Not Directly, Referenced | Monocytes (CD45): 30-F11, BioLegend  Monocytes (CD11b): M1/70, BioLegend  Monocytes (Ly6C): AL-21, BD Pharmingen | No | No | No | Not Directly, Referenced |
| Li et al., 2019 (67) | Monocytes | MoFlo (Beckman Coulter) | N/R | Yes | Monocytes (CD45): 30-F11, BioLegend  Monocytes (CD11b): M1/70, BioLegend | Yes | Yes | Yes | Yes |
| Hsieh et al., 2013 (68) | Bulk Macrophages | FACSAria (BD Biosciences) | FlowJo (Tree Star) | P/D | Macrophages (CD45): Ly5, eBioscience  Macrophages (CD11b): M1/70, Invitrogen or eBioscience | No | No | Yes | No |
| Hsieh et al., 2014 (69) | Bulk Macrophages | FACSAria (BD Biosciences) | FlowJo 9.6 (Tree Star) | P/D | Macrophages (CD45): Ly5, eBioscience  Macrophages (CD11b): M1/70, Invitrogen or eBioscience | No | No | No | No |
| DePaula-Silva et al. (70) | Bulk Macrophages | LSRFortessa X-20 Cell Analyzer (BD Biosciences) | FlowJo (Tree Star) | P/D | N/R | No | No | No | Yes |
| Lehmann et al. (71) | Bulk Macrophages | MoFlo Astrios (Beckman Coulter) | FlowJo (Tree Star) | Yes | N/R | Yes | Yes | Yes | Yes |
| Cai et al. (72) | Bulk Macrophages | FACSAria (BD Biosciences) | FlowJo X 10.07r2 | Yes | N/R | No | No | Yes | Yes |
| Martin et al. (73) | TMEM119^+^ Microglia | BD FACSVERSE (BD Biosciences) | FlowJo (FlowJo LLC) | Yes | N/R | Yes | Yes | No | Yes |
| Honarpisheh et al. (74) | TMEM119^+^ Microglia | Cytoflex-S (Beckman Coulter);  BD FACSMelody | FlowJo (Tree Star) | Yes | Microglia (CD11b): M1/70, BioLegend  Microglia (TMEM119): V3RT1GOsz, eBioscience  Microglia (P2RY12): S16007D (BioLegend) | Yes | Yes | Yes | Yes |
| Nirwane et al. (75) | TMEM119^+^ Microglia | ImageStreamX Mark II (Luminex) | IDEAS (Millipore) | Yes | Microglia (CD11b): M1/70, BioLegend  Microglia (CD45): 30-F11, BioLegend  Microglia (TMEM119): 28-3, Abcam | No | No | No | Yes |
| Spiteri et al. (76) | TMEM119^+^ Microglia | LSR II FACS (Becton Dickinson);  5-laser Spectral Aurora (Cytek Biosciences) | FACSDiva;  FlowJo v10.5 (Tree Star) | Yes | Microglia (CD45): 30-F11, BioLegend  Microglia (CD11b): M1/70, BioLegend and BD Biosciences  Microglia (CX3CR1): SA011F11, B, BioLegend  Microglia (P2RY12): S16007D, BioLegend | No | Yes | Yes | Yes |
| Perez-de-Puig et al. (77) | Neutrophils | FACS Canto II (Becton Dickinson) | FACSDiva (BD Biosciences); FlowJo v.7.6.5 (Tree Star) | Yes | Neutrophils (CD11b): M1/70, BD Pharmingen  Neutrophils (CD45): 30-F11, BD Pharmingen  Neutrophils (Ly6G): 1A8, BD Pharmingen | No | No | No | Yes |
| Garcia-Bonilla et al., 2014 (78) | Neutrophils | Accuri C6 (BD Biosciences); FACSVantage (BD Biosciences) | N/R | Yes | Neutrophils (CD11b): M1/70, BioLegend  Neutrophils (CD45): 30-F11, BioLegend  Neutrophils (Gr1): RB6-865, BioLegend | No | No | No | No |
| Makinde et al. (79) | Neutrophils | LSR II (BD Biosciences) | FlowJo (Tree Star) | Yes | Neutrophils (CD11b): M1/70, eBioscience  Neutrophils (CD45): 30-F11, BD Bioscience  Neutrophils (Ly6G): 1A8, BD Bioscience | No | No | No | No |
| Garcia-Bonilla et al., 2015 (80) | Neutrophils | Accuri C6 (BD Biosciences); FACSVantage (BD Biosciences) | N/R | P/D | Neutrophils (CD11b): M1/70, BioLegend  Neutrophils (CD45): 30-F11, BioLegend  Neutrophils (Ly6G): 1A8, BioLegend | No | No | No | No |
| Yao et al. (81) | Neutrophils | LSRII (BD Biosciences) | FlowJo v10 (TreeStar) | Yes | N/R | No | No | No | No |
| Posel et al. (82) | Neutrophils | FACS Canto II (BD Biosciences) | N/R | Yes | Neutrophils (CD45): 104  Neutrophils (Ly6G): 1A8 | Yes | No | No | Yes |
| Garcia-Culebras et al. (83) | Neutrophils | FACS Aria III; FACS Calibur (BD Biosciences) | Summit (Dako); FlowJo | P/D | Neutrophils (CD11b): M1/70, BioLegend  Neutrophils (CD45): 30-F11, BioLegend  Neutrophils (Ly6G): 1A8, BioLegend | No | No | No | No |
| Roy-O’Reily et al. (84) | Neutrophils | Cytoflex S Flow Cytometer (Beckmann Coluter) | FlowJo (TreeStar) | Yes | N/R | No | No | No | No |
|  | N/R: Not Reported  P/D: Partially Demonstrated | | | | | | | | |

**Suppl. Table 3. Mean values of wild-type/control mouse brain immune cell counts detected by flow cytometry (regardless of age and sex; standardized to 1 x 10^5^ total cells collected per sample)**

| **Suppl. Table 3. Mean values of wild-Type/Control Mouse Brain Immune Cell Counts Detected by Flow Cytometry (Regardless of Age and Gender; Standardized to 1 x 10^5^ Total Cells Collected Per Sample^1^)** | | | | | |
| --- | --- | --- | --- | --- | --- |
| **Cell Type** | **Raw Total Immune Cell Subset Count Reported Combined Mean (± SD)** | **Raw Total Overall Cell Count Collected Per Sample Reported Combined Mean (± SD)** | **Standardized Total Immune Cell Subset Count^1^ (± SD)** | **Standardized Total Immune Cell Subset % in Brain^2^(± SD)** | **References** |
| **Bulk CD4^+^ T cells** | 3,041 ± 3,428; n = 7 | 32,500 ± 24,749; n = 2 | 6,414 ± 6,148 | 6.41% ± 6.15 | Williams et al. (27)  Ferretti et al. (28)  Ito et al. (29)  Daglas et al. (30)  Pasciuto et al. (31)  Herz et al. (32)  Chu et al., 2014 (33) |
| **Bulk CD8^+^ T cells** | 2,383 ± 3,106; n = 7 | 66,000 ± 41,921; n = 4 | 4,093 ± 5,336 | 4.09% ± 5.34 | Daglas et al.(30)  Williams et al. (27)  Zhou et al. (34)  Shaw et al. (35)  Jurado et al. (36)  Unger et al. (37)  Ferretti et al. (28) |
| **DN T cells** | 181 ± 161; n = 2 | 12,500 ± 3,536; n = 2 | 1,816 ± 1,573 | 1.82% ± 1.57 | Chu et al., 2014 (33)  Meng et al. (38) |
| **T_REG_ T cells** | 630 ± 1,088; n = 4 | 106,000 ± 132,936; n = 2 | 863 ± 1,329 | 0.86% ± 1.33 | Ito et al. (29)  Beckmann et al. (39)  Li et al., 2021 (40)  Chen et al. (41)  Maes et al. (42) |
| **T_FH_ T cells** | 8 ± 0; n = 1 | N/R (Extrapolated)  84,000 ± 0; n =1 | 8 ± 0 | 0.008% ± 0 | Rayasam et al. (43) |
| **T_H1_ T cells** | 1,096 ± 1,394; n = 2 | 62,500 ± 53,033; n = 2 | 2,392 ± 2,704 | 2.39% ± 2.70 | Williams et al. (27)  Li et al., 2021 (40) |
| **T_H17_ T cells** | 3,150 ± 636; n = 2 | N/R (Extrapolated)  1,000,000 ± 0; n = 2 | 315 ± 67 | 0.32% ± 0.06 | Guo et al. (44)  Beurel et al. (45) |
| **T_EM_ Bulk CD4^+^ T cells** | 503 ± 0; n = 1 | 50,000 ± 0; n =1 | 1,006 ± 0 | 1.01% ± 0 | Daglas et al. (30) |
| **T_EM_ Bulk CD8^+^ T cells** | 377 ± 0; n = 1 | 50,000 ± 0; n =1 | 754 ± 0 | 0.75% ± 0 | Daglas et al. (30) |
| **NK cells** | 27,982 ± 44,684; n = 7 | 1,000,000 ± 0; n = 1 | 1,278 ± 2,643 | 1.28% ± 2.64 | He et al. (46)  Jin et al. (47)  Zhang et al. (48)  Li et al., 2020 (49)  D’Alessandro et al. (50)  Chu et al., 2014 (33)  Earls et al. (51) |
| **DCs** | 895 ± 1,331; n = 5 | 3,061,250 ± 4,311,584; n = 2 | 41 ± 48 | 0.04% ± 0.05 | Dando et al. (52)  Clarkson et al. (53)  Gelderblom et al. (54)  Gobel et al. (54)  Chu et al., 2014 (33) |
| **B cells** | 6,748 ± 18,277; n = 6 | 1,000,000 ± 0; n = 2 | 1,285 ± 2,540 | 1.28% ± 2.54 | Brioschi et al. (56)  Chu et al., 2014 (33)  Pierson et al. (57)  Korin et al. (58)  Liu et al. (59)  Mracsko et al. (60) |
| **Monocytes** | 31,683 ± 69,662; n = 7 | 2,142,500 ± 1,212,688; n = 2 | 891 ± 2,118 | 0.89% ± 2.11 | Trahanas et al. (61)  Miro-Mur et al. (62)  Ritzel et al. (63)  Cazareth et al. (64)  Chu et al., 2015 (65)  Peralta et al. (66)  Li et al., 2019 (67) |
| **Bulk Macrophages** | 2,634 ± 2,196; n = 8 | 650,000 ± 494,975; n = 2 | 327 ± 412 | 0.33% ± 0.41 | Hsieh et al., 2013 (68)  Cazareth et al. (64)  Hsieh et al., 2014 (69)  Chu et al., 2015 (65)  DePaula-Silva et al. (70)  Lehmann et al. (71)  Cai et al. (72) |
| **TMEM119^+^ Microglia** | 90,323 ± 104,555; n = 4 | 316,667 ± 315,331; n = 3 | 28,520 ± 33,017 | 28.5% ± 33.0 | Martin et al. (73)  Honarpisheh et al. (74)  Nirwane et al. (75)  Spiteri et al. (76) |
| **Neutrophils** | 1,821 ± 3,624; n = 9 | 810,000 ± 438,406; n = 2 | 193 ± 250 | 0.19% ± 0.25 | Perez-de-Puig et al. (77)  Garcia-Bonilla et al., 2014 (78)  Makinde et al. (79)  Garcia-Bonilla et al., 2015 (80)  He et al. (46)  Yao et al. (81)  Posel et al. (82)  Garcia-Culebras et al. (83)  Roy-O’Reily et al. (84) |
| ^1^: $\frac{a}{b}=\frac{X}{{10}^{5} Total Cells Collected}$; a: Raw Total Immune Cell Subset Count; b: Raw Total Overall Cell Count Collected Per Sample; X: Standardized Total Immune Cell Subset Count  ^2^: ([Standardized Total Immune Cell Subset Count/10^5^ Total Cells Collected] *100)  n: Number of Studies Data Derived From  The following subsets were not documented in any study: T_H2_ T cells; Naïve-Like Bulk CD4^+^ T cells; Naïve-Like Bulk CD8^+^ T cells; Naïve-Like T_REG_ T cells; Naïve-Like T_FH_ T cells; Naïve-Like T_H1_ T cells; Naïve-Like T_H2_ T cells; Naïve-Like T_H17_ T cells; T_CM_ Bulk CD4^+^ T cells; T_CM_ Bulk CD8^+^ T cells; T_CM_ T_REG_ T cells; T_CM_ T_FH_ T cells; T_CM_ T_H1_ T cells; T_CM_ T_H2_ T cells; T_CM_ T_H17_ T cells; T_EM_ T_REG_ T cells; T_EM_ T_FH_ T cells; T_EM_ T_H1_ T cells; T_EM_ T_H2_ T cells; T_EM_ T_H17_ T cells; T_EMRA_-Like Bulk CD4^+^ T cells; T_EMRA_-Like Bulk CD8^+^ T cells; T_EMRA_-Like T_REG_ T cells; T_EMRA_-Like T_FH_ T cells; T_EMRA_-Like T_H1_ T cells; T_EMRA_-Like T_H2_ T cells; T_EMRA_-Like T_H17_ T cells; M1 Macrophages; M2 Macrophages. | | | | | |
